# Spatial modelling improves genetic evaluation in smallholder breeding programs

**DOI:** 10.1101/2020.06.01.128868

**Authors:** Maria L. Selle, Ingelin Steinsland, Owen Powell, John M. Hickey, Gregor Gorjanc

## Abstract

Breeders and geneticists use statistical models for genetic evaluation of animals to separate genetic and environmental effects on phenotype. A common way to separate these effects is to model a descriptor of an environment, a contemporary group or herd, and account for genetic relationship between animals across the environments. However, separating the genetic and environmental effects in smallholder systems is challenging due to small herd sizes and weak genetic connectedness across herds. Our hypothesis was that accounting for spatial relationships between nearby herds can improve genetic evaluation in smallholder systems. Further, geographically referenced environmental covariates are increasingly available and could be used to model underlying sources of the spatial relationships. The objective of this study was therefore to evaluate the potential of spatial modelling to improve genetic evaluation in smallholder systems. We focus solely on dairy cattle smallholder systems.

We performed simulations and real dairy cattle data analysis to test our hypothesis. We used a range of models to account for environmental variation by estimating herd and spatial effects. We compared these models using pedigree or genomic data.

The results show that in smallholder systems (i) standard models are not able to separate genetic and environmental effects, (ii) spatial modelling increases accuracy of genetic evaluation for phenotyped and non-phenotyped animals, (iii) environmental covariates do not substantially improve accuracy of genetic evaluation beyond simple distance-driven spatial relationships between herds, (iv) the benefit of spatial modelling was the largest when the genetic and environmental effects were hard to separate and (v) spatial modelling was beneficial when using either pedigree or genomic data.

We have demonstrated the potential of spatial modelling to improve genetic evaluation in smallholder systems. This improvement is driven by establishing environmental connectedness between herds that enhances separation of the genetic and environmental effects. We suggest routine spatial modelling in genetic evaluations, particularly for smallholder systems. Spatial modelling could also have major impact in studies of human and wild populations.

## Background

This work evaluates the potential of spatial modelling to improve genetic evaluation of animals in smallholder systems. Over the past century genetic selection of dairy cattle has had a big impact on the increase in milk production in developed countries (Weigel et al., 2017). For example, the average milk production of US Holstein cows has almost doubled between 1960 and 2000, and more than half of this is due to genetic improvement (Dekkers and Hospital, 2002). However, the same improvement in livestock productivity has not been achieved in low to middle income countries, for example in East Africa. For example, Rademaker et al. (2016) reported that milk yields of smallholder producers in Kenya are about 5–8 litres per cow per day, which is several-fold smaller than milk yields of large-scale commercial farmers around the world. Milk yields in low to middle income countries are constrained by environmental, technological and infrastructural difficulties (Philipsson et al., 2011; Majiwa et al., 2017). Whereas the large-scale commercial farmers measure phenotypes accurately, keep records of performance and pedigree, the smallholders usually do not keep records and the absence of routine phenotyping systems reduces accuracy of these records (Ojango et al., 2019; Powell et al., 2019).

To perform accurate genetic evaluation of animals in a breeding program, a sufficient amount of data is needed, and the data should be properly structured (Foulley et al., 1990; Jorjani et al., 2001; Powell et al., 2019). In developed countries, a small number of large-scale commercial farms produce most of milk and there is widespread use of artificial insemination that establishes strong genetic connectedness between herds. On the other hand, in many smallholder systems the smallholder farms contribute significantly to milk production and there is low usage of artificial insemination with consequent weak genetic connectedness between herds. For example, smallholder milk-producing households in Kenya, who own one to three cows, represent approximately 80% of the national dairy herds (Rademaker et al., 2016). Further, 87% of Kenyan farmers asked in a survey used natural mating services rather than artificial insemination, even though 54% reported they preferred artificial insemination (Lawrence et al., 2015). Similar values for the proportion of natural mating and artificial insemination was reported by Bebe et al. (2003); Baltenweck et al. (2004).

Small herd sizes and weak genetic connectedness across herds challenge accurate genetic evaluation because it is hard to separate the genetic and environmental effects. When the herd sizes are small, for example if a herd consists of only one cow, it is not at all possible to separate the genetic and environmental effects on the phenotype. When the genetic connectedness is weak, the genetic relationship between animals in different herds is weak, which also makes it hard to share information between animals and herds to improve separation of genetic and environmental effects. Since most smallholders mate cows with their own or neighbour’s bull, it is reasonable to assume that most farmers close in distance, for example farmers belonging to the same village, use the same bulls. This creates genetic connectedness across herds close in distance even though the overall genetic connectedness across the country is weak.

In the statistical models for genetic evaluations, the genetic effect is modelled using expected or realised genetic relationship between animals, respectively derived from pedigree or genomic data. A herd effect, or a herd-year-season effect is often included as the main environmental effect (Henderson et al., 1984; Visscher and Goddard, 1993; Ojango et al., 2019; Pereira et al., 2019; Mrode, 2014). When herd sizes are small the herds are treated as random, as this has been found to give higher accuracy than treating them as fixed (Visscher and Goddard, 1993; Frey et al., 1997; Schaeffer, 2018; Powell et al., 2019). In the extreme case of single animal per herd, modelling herds as random is in fact the only possible approach (Powell et al., 2019). In addition, including other factors and covariates in the statistical models is a way of including information in the model that can further enhance the separation of genetic and environmental effects.

Environmental effects can be on management (herd) level, or on a larger scale, likely shared by herds close in distance. Examples of environmental effects on management level are education, age, experience, land size, use of natural mating or artificial insemination. Some of these effects can be similar for herds close in distance. The quality of feeding used in farms is likely similar in farms belonging to the same villages, and vaccination in farms is likely correlated with local, regional or national government policies. These will be similar between herds close in distance, for example herds that belong to the same village. Examples of large scale environmental effects are climate effects, proximity to roads, markets, and towns, and government policies. Many of the environmental effects, both the ones on management level and the large scale effects, can be assumed to be spatially correlated. We will refer to environmental effects on management level as herd effects, and large scale environmental effects as spatial effects.

There are multiple spatial models that could be used in an animal breeding context. A prerequisite for this is that collected data are geographically referenced, meaning their geographical location in country must be known. Geographical location can be described coarsely with regions, such as administrative counties or districts, or precisely with point coordinates. For an application of region-based models in an animal breeding context see Sæbø and Frigessi (2004), who modelled veterinary district as an environmental effect with relationships between neighbouring districts following the so called CAR model of Besag (1974); Rue and Held (2005). We focus on coordinate-based models (often referred to as geostatistical models (Rue and Held, 2005; Gelfand et al., 2010; Cressie, 2015; Cressie and Wikle, 2015)) to account for fine-grained spatial relationships between smallholder farms. The only requirement for a coordinate-based models is that we collect herd coordinates and then all data pertaining to a herd is point-referenced. For a herd *i* we define a tuple ***w**_i_* that typically contains two-dimensional coordinates (latitude and longitude), but further extensions are possible (Lindgren et al., 2011; Ingebrigtsen et al., 2014). The observation at specific locations and locations themselves can vary continuously over a geographical region. A common model for such continuous spatial processes is a Gaussian random field where we model observations at a set of locations (*y*(***w***_1_), …, *y*(***w**_n_*)) with a multivariate normal distribution with mean ***μ*** and a spatially structured covariance matrix **Σ** (Rue and Held, 2005). The same approach can also be used as a model component in the context of a linear mixed model (Rue and Held, 2005), as is the case with genetic effects, but in the spatial context we account for relationships between locations. There are multiple possible covariance functions for spatial modelling. Most of them assume stationarity and isotropy so that ***μ***(***w***) = ***μ*** and spatial covariance between locations is a function of Euclidian distance between locations and model parameters, such as variance. The most commonly used is the Matérn covariance function (Matérn, 1960).

Modelling with continuously indexed Gaussian random fields is computationally challenging because they give rise to dense precision (covariance inverse) matrices that are numerically expensive to factorise (Rue and Held, 2005), as is the case with genomic models VanRaden (2008); Gorjanc et al. (2018). Gaussian Markov random fields approximate Gaussian random fields by inducing conditional independence (Markov) assumptions, which increases sparsity of the precision matrix and reduces computational complexity. Lindgren et al. (2011) showed how to construct an explicit link between (some) Gaussian random fields and Gaussian Markov random fields via solution of stochastic partial differential equations. They also proposed use of finite element methods to further reduce computational complexity. This approach allows implementation of computationally efficient numerical methods for spatial modelling of large scale point-referenced data. Assuming conditional independence to enable scalable computations has also been recently proposed for dense genomic models (Misztal et al., 2014; Misztal, 2016).

The aim of this study was to evaluate the potential of spatial modelling in addition to modelling independent herd effects to improve genetic evaluation in smallholder systems, and to determine if the impact was dependent on the genetic connectedness across the herds, and the use of pedigree or genomic data for modelling genetic relationships. In addition we wanted to test whether adding environmental covariates was beneficial beyond the simple distance-driven spatial relationships between herds.

We performed a simulation study that resembled smallholder systems commonly observed in East Africa with small herd sizes. We evaluated scenarios with different genetic connectedness across herds, herd distribution and spatial variation. The results showed that spatial modelling improved genetic evaluations, especially with weak genetic connectedness. We also analysed real data from a dairy cattle programme and the results indicated that the standard and spatial models separated the genetic and environmental effects in different ways.

## Material and methods

We first introduce the data used in the analyses; a simulated smallholder dairy cattle data, and a real cattle data. Then we present the statistical models used for genetic evaluation and how we fitted and evaluated the models. Scripts for data simulation and model fitting are available in [Additional file 2].

### Simulation

We first used simulation to evaluate the potential of spatial modelling to improve genetic evaluation. The simulated data resembled the smallholder systems commonly observed in East Africa with small herds clustered in villages and varying level of genetic connectedness. In summary, we simulated phenotype observations *y_i_* as:

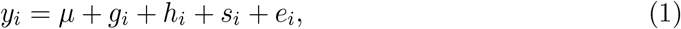

where *μ* is population mean, *g_i_* is the additive genetic effect of individual *i*, 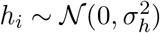 is the herd effect with 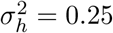, *s_i_* is the spatial effect, and 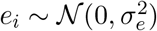 is an independent residual effect with 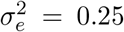. In Figure 1, we show a conceptual illustration of the simulation. The top left panel shows the phenotypes, and the remaining panels show the genetic, herd and spatial effects, while we omitted the residuals from the illustration. Please note the most bottom-right village (cluster) with high genetic merit animals, but intermediate phenotypes due to spatial effects. We describe in detail how the genetic and spatial effects were generated below.

**Figure 1:**
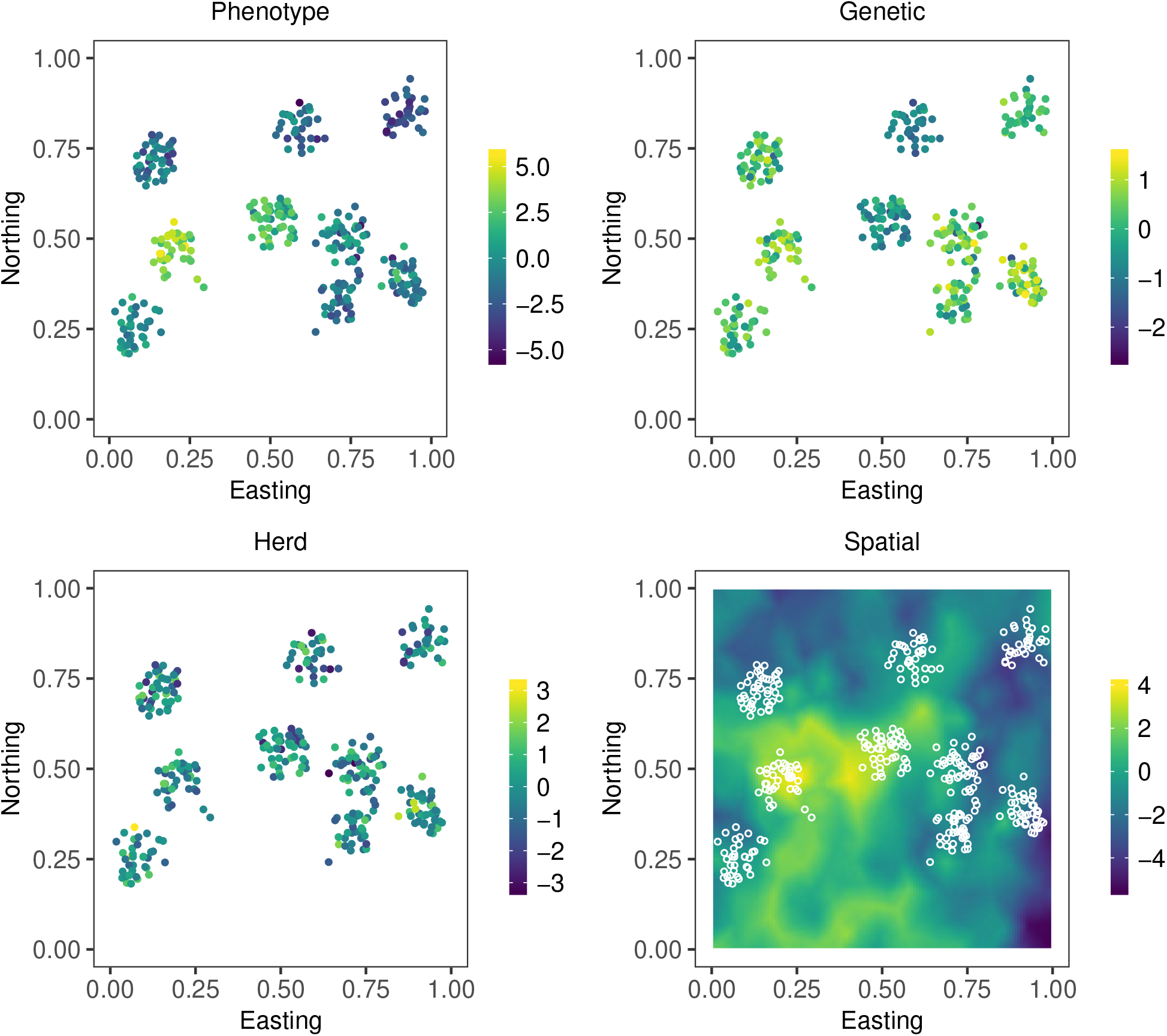
Illustration of the simulation. Each point denotes an animal, their location in a country and colour of the point denotes value of phenotype and underlying genetic, herd and spatial effects (residual not shown)

We simulated the data under three scenarios of genetic connectedness, from weak genetic connectedness between herds from different villages to strong genetic connectedness across all herds regardless of village. We generated 60 independent data sets for each scenario of genetic connectedness.

### Simulation of founders

We first simulated a genome consisting of 10 chromosome pairs with cattle genomic and demographic parameters (MacLeod et al., 2013). To this end we used the Markovian Coalescent Simulator (Chen et al., 2009) and AlphaSimR (Faux et al., 2016; Gaynor et al., 2019) to simulate genome sequences for 5,000 founder individuals, who served as the initial parents. For each chromosome, we randomly chose segregating sites in the founders’ sequences to serve as 5,000 single-nucleotide polymorphism (SNP) markers and 1,000 quantitative trait loci (QTL) per chromosome, yielding in total 50,000 SNPs and 10,000 QTL.

Then we simulated a single complex trait with additive architecture by sampling QTL allele substitution effects from a standard normal distribution. We multiplied these with individuals’ QTLs and summed them into the true breeding value. Then we simulated phenotypes with different heritabilities for cows (*h*^2^ = 0.3) and bulls (*h*^2^ = 0.8) to reflect different amounts of information per gender. These phenotypes were used for the initial assignment of bulls and their selection throughout the evaluation phase.

### Population simulation

We created 100 villages, each consisting of 20 herds, with herd sizes generated from a zero truncated Poisson distribution with parameter λ = 1.5. The 110 best males from the founder individuals (based on true genetic values) were assigned as breeding bulls, 100 of them as natural mating bulls, and 10 of them as artificial insemination bulls. The remaining founders were considered as cows, and were randomly placed in the herds according to their predetermined size. Since the herd sizes were sampled, we did not have the same number of individuals in each independent replicate. On average there were 3,860 cows in total, and the cows not assigned to a herd were discarded.

We positioned the 100 villages by assuming a square country and sampling village coordinates in north-south direction and in east-west direction from a uniform distribution on (0,1). We then positioned the 2,000 herds by sampling their coordinates 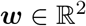 from a bi-variate normal distribution with mean from the corresponding village coordinates and location variance 3.5 · 10^−4^***I***_2×2_. This clustered the herds into villages. We chose the location variance to achieve reasonable spread and clustering. We tested the sensitivity of results to this simulation parameter.

We created three different levels of genetic connectedness by controlling the breeding strategy. To achieve weak genetic connectedness each village used their own bull, meaning that the cows were strongly related within the village and nominally unrelated across villages. However, there was still some base level genetic relationship due to shared population history. To achieve intermediate genetic connectedness each village used their own bull for mating in 75% of the herds, while the remaining herds in the village used one of the 10 artificial insemination bulls at random, meaning that cows were still strongly related within villages, and somewhat related across villages. To achieve strong genetic connectedness the 100 natural mating bulls were randomly mated to cows across all herds and villages, meaning that cows were equally related within and across villages. For this last scenario we used the 100 natural mating bulls instead of the 10 artificial insemination bulls in order to maintain relatively high degree of genetic diversity and with this more challenging situation for separation of environmental and genetic effects.

The three scenarios were then simulated over twelve discrete generations of breeding. Within each farm, we replaced old cows by newborn female calves. The cows who had male calves were not replaced, and their calves were evaluated on suitability as natural mating bulls if they came from a farm using natural mating bulls, or as artificial insemination bulls if they came from a farm using artificial insemination.

In the 11th generation we scaled the true breeding values to have mean 0 and variance 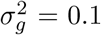 and used them as genetic effects in the model for phenotype observation *y_i_* (1) with 3,860 records on average. In addition the female calves in the 12th generation were kept for prediction purposes. To ease the computations with the genome based model, we predicted breeding values for randomly chosen 200 calves in the 12th generation.

### Simulation of spatial effects

We simulated spatial effects from multiple Gaussian random fields to mimic several sources of environmental effects, both on large and small scale. We imagined that these different sources could be temperature, precipitation, elevation, land size, proximity to markets and towns, availability of extension services, vaccine use, local and regional policies etc. We simulated the effects of 8 such processes ***v***_*k*_, *k* = 1,…, 8 at the herd locations from a Gaussian random field with mean 0 and a Matérn covariance function (Matérn, 1960). The Matérn covariance function between locations ***w**_i_*, 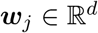 is:

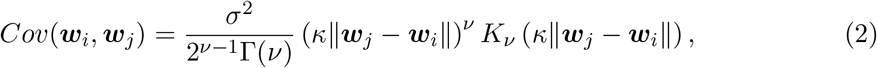

where *K_v_* is the modified Bessel function of the second kind and the order *v* > 0 determines the mean-square differentiability of the field. The parameter *κ* can be expressed as 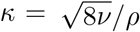, where *ρ* > 0 is the range parameter describing the distance where correlation between two points is near 0.1, and *σ*^2^ is the marginal variance. We varied these parameters to simulate processes on large and small scale and with different properties. Specifically, we sampled the range parameter *ρ* for each of the processes ***v**_k_* from a uniform distribution on (0.1,0.5), set the marginal variance *σ*^2^ to either 0.2 or 0.3 with equal probability, and fixed the parameter *v* to 1.

We finally summed the 8 processes to obtain the total spatial effect (Figure 1) for all herd locations ***s***, with ***s***(***w**_i_*) being the total spatial effect at location ***w**_i_*. We differentially emphasised some processes according to:

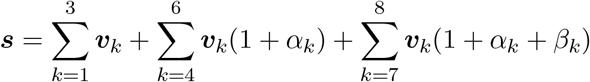

with the weights *α*, *β* ~ Uniform(−0.5,0.5). We scaled the spatial effects to have mean 0 and variance 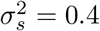.

### Environmental covariates

Further we assumed that some spatial processes can be observed as environmental covariates at herd locations, possibly with some noise. We took the 8 real processes and sampled two more (with mean 0 and a Matérn covariance function), but these two did not affect the phenotype.

For the spatial processes ***v***_1_, ***v***_2_, and ***v***_3_ we assumed that we could observe the spatial covariates perfectly without error, which could be reasonable for some covariate effects like average temperature and precipitation.

For the spatial processes ***v***_4_, ***v***_5_, and ***v***_6_ we assumed that they were harder to observe accurately, so we added normal distributed error terms with mean 0 and variance equal to 10% of the process marginal variance. This could be reasonable for some covariates that are difficult to measure or that vary with time, it could for example be difficult to quantify the amount of different types of feeding.

For the spatial processes ***v***_7_ and ***v***_8_ we assumed that we could only observe categorical realisations of the continuous effects, for example distance to markets and towns could be categorised as either a rural or urban area. For the process ***v***_7_ we created a two-level categorical covariate by sampling a threshold from a uniform distribution between one standard deviation from the mean of ***v***_7_ in both negative and positive direction. Values of ***v***_7_ above the threshold were assigned one level, and values below were assigned the other level. For the process ***v***_8_ we created a three-level categorical covariate by sampling two thresholds. The lower threshold was sampled from a uniform distribution between two standard deviations below the mean of ***v***_8_ and the mean of ***v***_8_. The upper threshold was sampled from a uniform distribution between the mean of ***v***_8_ and two standard deviations above the mean of ***v***_8_. The values of ***v***_8_ were then assigned one of three categorical levels depending on which thresholds they were between.

### Changing the proportion of spatial variance and herd clustering

To evaluate how the models performed when there was no or little spatial effect on the phenotype, we created scenarios with different proportions of spatial variance relative to the sum of herd effect variance and spatial variance so that the total variation between herds was constant. We kept 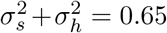, and let 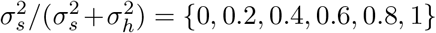. This was repeated for 30 of the data sets.

We also evaluated the importance of how closely the herds were clustered around villages. To do this we varied the location variance of the bi-variate distribution for the herd coordinates 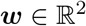 from 1.0·10^−4^***I***_2×2_ (strong clustering), 3.5·10^−4^***I***_2×2_ (intermediate clustering) to 9.0 · 10^−4^***I***_2×2_ (weak clustering). This was repeated for each of the 60 data sets.

### Real cattle data

We then analysed phenotypic data for 30,314 Brown-Swiss cattle data from Slovenia collected between 2004 and 2019, from 2,012 different herds. The data included a trait describing a body confirmation measure, year and scorer of the data, cattle age, stage of lactation, year and month of calving, and herd location coordinates. In addition the data contained a pedigree for 56,465 animals including the cows with phenotypes. We analysed the trait, which was standardised by subtracting the phenotypic mean and dividing by the phenotypic standard deviation.

The average herd size was approximately 15 cows per herd, and most cows belonged to herds consisting of more than five animals. To imitate data similar to the typical smallholder system, with few individuals per herd, we used a subset of the full data. We sampled 3,800 individuals without replacement, with sampling probability equal to the inverse herd size, meaning that larger herds had fewer records in the data subset. The subset contained cows from 1,838 different herds, and the average herd size was about 2 cows per herd. The herds were spread over most of Slovenia, and their locations are shown in [Additional file 1, Figure 2].

**Figure 2:**
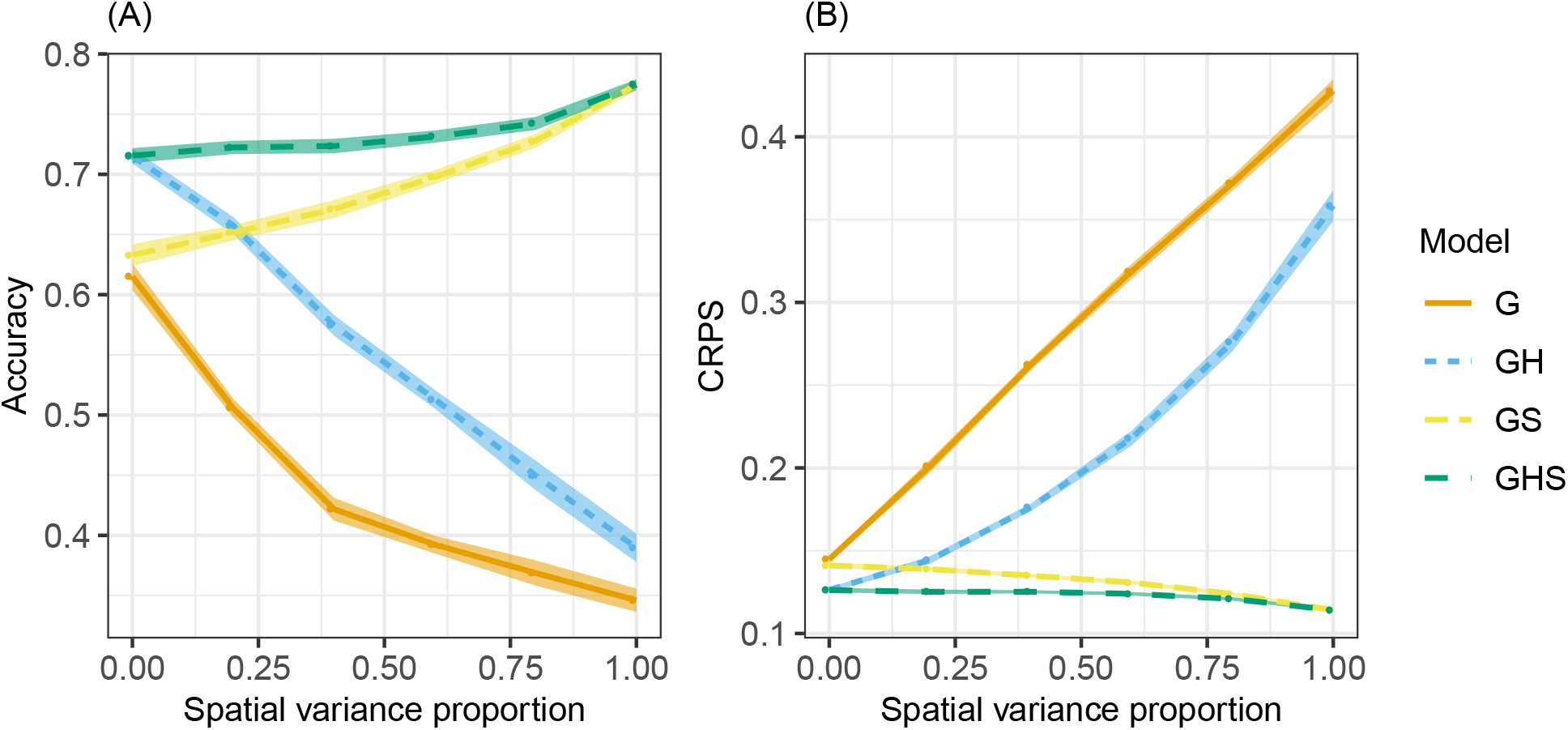
Average accuracy (A) and CRPS (smaller is better) (B) with 95% confidence intervals for estimated breeding values by proportion of spatial variance in the sum of spatial and herd variance in scenario with intermediate genetic connectedness and using genomic model

### Statistical models

The following model was fitted to the observed phenotype *y_i_* of individual *i* = 1, …, *n*:

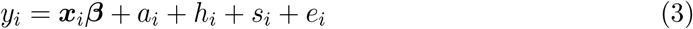

where ***β*** is a vector of fixed covariate effects, including a common intercept, with known covariate vector *x_i_* and 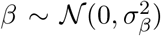, *a_i_* is the additive genetic effect (breeding value), *h_i_* is the herd effect with 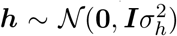, *s_i_* is the spatial effect for the herd at location 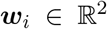 modelled with a Gaussian random field with ***μ*** = **0** and Matérn covariance function as given in (2), and *e_i_* is a residual effect with 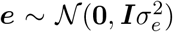. Albeit the data generation model and this statistical model are similar we note that the statistical model is not ”aware” of the 10,000 true QTL affects and 8 true spatial processes.

We modelled the genetic effect (breeding value) using relationship matrix based either on pedigree or genome data. For the pedigree based model we assumed 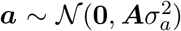, where ***A*** is the pedigree relationship matrix (Lynch et al., 1998). We used pedigree for the phenotyped individuals (11th generation), their offspring (12th generation), and three previous generations (8-10th). For the genome based model we assumed 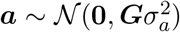, where ***G*** is the genomic relationship matrix calculated from ***G*** = ***ZZ**^T^*/*k*, ***Z*** was a column-centered SNP marker matrix, and *k* = 2∑_*lql*_(1 ‒ *q_l_*) with *q_l_* being allele frequency of marker *l* (VanRaden, 2008).

### Prior distributions for hyper-parameters

We used a full Bayesian analysis which requires prior distributions for all model parameters. For the intercept and fixed effects we assumed 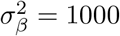, and for the remaining variance parameters and the spatial range we assumed penalised complexity priors (Simpson et al., 2017), which are proper priors that penalise model complexity to avoid over-fitting. The penalised complexity prior for variance parameters can be specified through a quantile *u* and a probability α which satisfy Prob(*σ* > *u*) = *α*, and the penalised complexity prior for the spatial range parameter through a quantile *u* and a probability *α* which satisfy Prob(*ρ* < *u*) = *α*. For the variances and spatial range we assumed penalised complexity prior distributions with quantiles u and probabilities *α*. We state the parameters *u* and *α* in Table 1.

**Table 1:** Parameters *u* and *α* for the penalised complexity priors of hyper-parameters by fitted models to the simulated^1^ and real^2^ data (see subsection Prior distributions for hyper-parameters)

| Model | $u_e, \alpha_e$ | $u_a, \alpha_a$ | $u_h, \alpha_h$ | $u_s, \alpha_s$ | $u_\rho, \alpha_\rho^1$ | $u_\rho, \alpha_\rho^2$ |
| --- | --- | --- | --- | --- | --- | --- |
| G | 0.30, 0.50 | 0.10, 0.50 | - | - | - | - |
| GH | 0.15, 0.50 | 0.10, 0.50 | 0.25, 0.50 | - | - | - |
| GS | 0.15, 0.50 | 0.10, 0.50 | - | 0.25, 0.50 | 0.60, 0.95 | 50, 0.80 |
| GHS | 0.15, 0.50 | 0.10, 0.50 | 0.15, 0.50 | 0.10, 0.50 | 0.60, 0.95 | 50, 0.80 |

### Fitted models to the simulation data

We fitted five different models to the simulated data: G, GH, GS, GHS and GHSC. All models had an intercept *β*_0_, a genetic effect *a_i_*, and a residual effect *e_i_*. G had no additional model components, GH had in addition a herd effect *h_i_*, GS had in addition a spatial effect *s_i_*, GHS had in addition both a herd effect and a spatial effect, and GHSC had in addition a herd effect, a spatial effect and the environmental covariates *z_i_*. The models are summarised as:

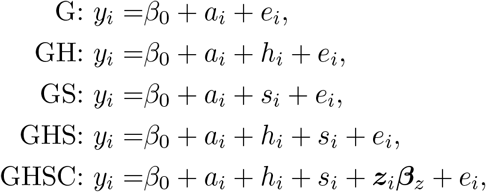

where ***z**_i_* is the vector of environmental covariates for individual *i* and 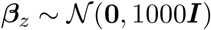 is a vector of environmental covariate effects. The other effects were distributed as described above for (3).

### Model evaluation with simulated data

We will refer to the mean posterior genetic effect for phenotyped individuals as the estimated breeding values, and the mean posterior genetic effect for non-phenotyped individuals as the predicted breeding values. We evaluated the models using correlation (accuracy) between the true and estimated or predicted breeding values, and the continuous rank probability score (CRPS) (Gneiting and Raftery, 2007), comparing the whole posterior distribution for breeding values to the true breeding values. The CRPS takes into account the whole posterior distribution, meaning it compares the estimated mean posterior value with the true value while taking into account the spread of the posterior distribution. The CRPS is negatively oriented, which means that lower CRPS values indicates a better estimate of breeding value.

### Fitted models to the real data

We fitted four different models to the real data that were structurally the same as models fitted to the simulated data: G, GH, GS, and GHS. The only difference was in fixed effects that are part of the routine genetic evaluation for the analysed trait and population; an intercept *β*_0_, three categorical fixed effects (one describing the year and scorer of the data, one describing cow age and stage of lactation, and one describing year and month of calving). The genetic effect was estimated using the available pedigree. For the variances and spatial range we assumed penalised complexity prior distributions with quantiles *u* and probabilities *α* shown in Table 1.

We used the deviance information criterion (DIC) (Spiegelhalter et al., 2002) to compare the model fit between the models fitted to the case study data. The DIC is widely used to compare model fit between different hierarchical Bayesian models while also assessing the model complexity. Lower values of the DIC indicate a better model fit.

### Inference

For inference, we used the Bayesian numerical approximation procedure known as the Integrated Nested Laplace Approximations (INLA) introduced by Rue et al. (2009), with further developments described in Martins et al. (2013) and implementation available in R-INLA package. INLA are suited for the class of latent/hierarchical Gaussian models, which includes for example generalised linear (mixed) models, generalised additive (mixed) models, spline smoothing methods, and the models used in this study. INLA calculate marginal posterior distributions for all model parameters (fixed and random effects, and hyper-parameters) and linear combinations of effects without using sampling-based methods such as Markov chain Monte Carlo (MCMC). For an in-depth description of INLA see Rue et al. (2009); Martins et al. (2013) and the recent review Rue et al. (2017).

## Results

In this section, we present the results from fitting the models to the simulated and real data. For simulation we compare accuracy and CRPS of the estimated and predicted breeding values for the tested models. For real data we present the posterior variances, the deviance information criterion, the estimated spatial effects, and how the estimated breeding values differ with and without spatial modelling. All results indicate that spatial modelling improves genetic evaluation.

### Simulated data

This section presents the results from the simulation study, where the models G, GH, GS, GHS and GHSC were fitted to data with different genetic connectedness. Overall, the results showed that in smallholder systems (1) spatial modelling increased accuracy of estimating and predicting breeding values, (2) the environmental covariates did not improve the accuracy substantially beyond the distance based spatial model, (3) the models without spatial effects were not able to separate genetic and spatial environmental effects, (4) the benefit of spatial modelling was largest when the genetic and environmental effects were strongly confounded, (5) spatial modelling in addition to the independent random herd effect did not decrease the accuracy even when there was no spatial effects, and (6) when environmental and genetic effects were confounded the accuracy improved when herds were weakly clustered rather than strongly clustered.

### Spatial modelling increases accuracy

Spatial modelling increased accuracy of estimated and predicted breeding values. Table 2 presents the accuracy for all models and genetic connectedness scenarios. Across all scenarios, model GHS gave the highest accuracy. The second best was GS, third was GH, and the worst was G. We expectedly see that genomic data improved the accuracy compared to using pedigree, and that the estimated breeding values were more accurate than the predicted. With weak genetic connectedness the accuracy was low and comparable for estimation and prediction, and the pedigree models had almost as high accuracy as the genomic models. This is reasonable since the animals were strongly related within the villages under this scenario.

**Table 2:** Average accuracy of estimated breeding values (EBV) and predicted breeding values (PBV) by genetic connectedness (weak, intermediate and strong) and model with intermediate clustering of herds. Standard error for most values had an order of magnitude 10^−3^ with few an order of magnitude 10^−2^.

|  | Weak |  | Intermediate |  | Strong |  |
| --- | --- | --- | --- | --- | --- | --- |
|  | EBV | PBV | EBV | PBV | EBV | PBV |
| **Pedigree** |  |  |  |  |  |  |
| G | 0.33 | 0.28 | 0.32 | 0.18 | 0.32 | 0.20 |
| GH | 0.36 | 0.29 | 0.41 | 0.22 | 0.42 | 0.25 |
| GS | 0.52 | 0.50 | 0.56 | 0.34 | 0.55 | 0.35 |
| GHS | 0.54 | 0.52 | 0.58 | 0.36 | 0.57 | 0.37 |
| GHSC | 0.57 | 0.55 | 0.59 | 0.36 | 0.58 | 0.37 |
| **Genomic** |  |  |  |  |  |  |
| G | 0.33 | 0.32 | 0.40 | 0.29 | 0.42 | 0.32 |
| GH | 0.36 | 0.33 | 0.51 | 0.38 | 0.59 | 0.46 |
| GS | 0.58 | 0.56 | 0.70 | 0.54 | 0.72 | 0.57 |
| GHS | 0.63 | 0.60 | 0.74 | 0.57 | 0.75 | 0.60 |
| GHSC | 0.64 | 0.62 | 0.74 | 0.58 | 0.75 | 0.60 |

Table 3 presents the average CRPS. The trends in the CRPS were the same as for the accuracy, with GHS having the lowest (best) CRPS. Again, we expectedly see that genomic data improved the CRPS compared to using pedigree, and in most cases average CRPS for the estimation were lower than for the prediction, but in some cases the average CRPS for prediction were slightly lower than for estimation. This improved CRPS for prediction was observed for models that did not model environmental variation and had lower accuracy, so the lower (better) CRPS indicates that those models underestimated prediction uncertainty.

**Table 3:**
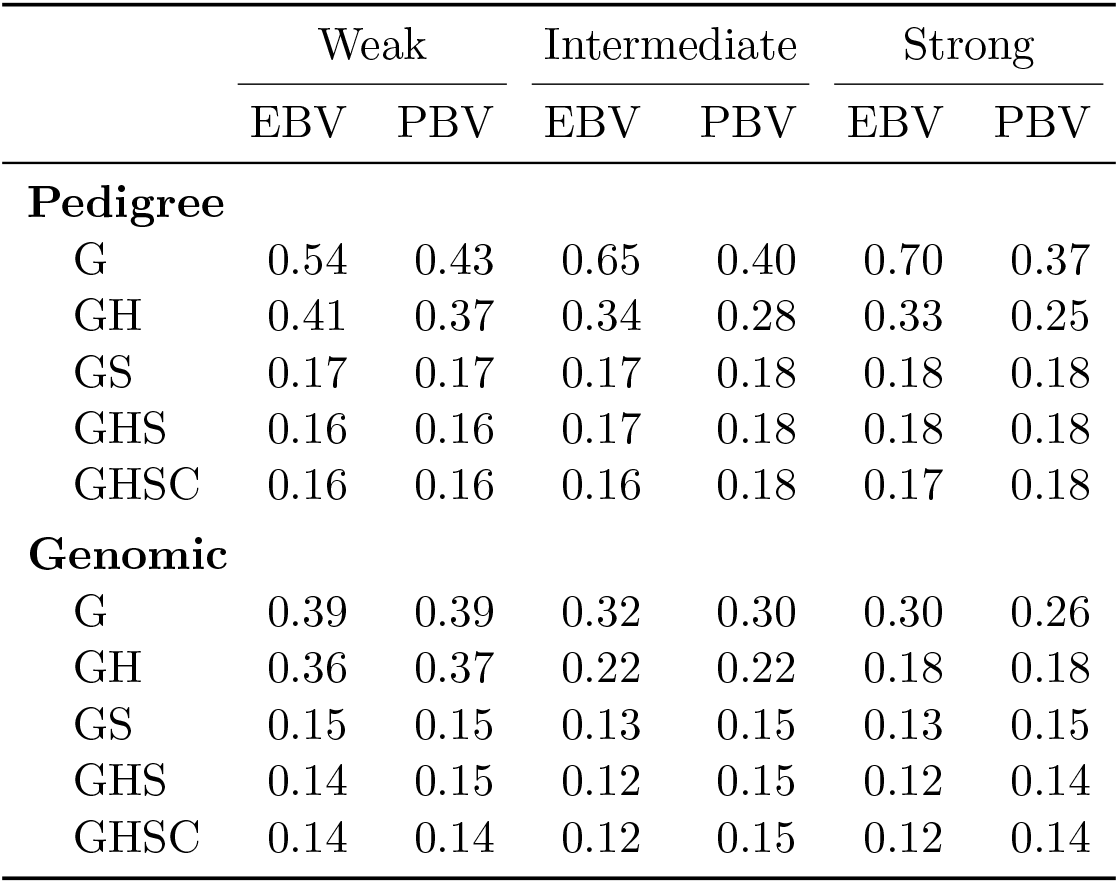
Average CRPS of estimated breeding values (EBV) and predicted breeding values (PBV) by genetic connectedness (weak, intermediate and strong) and model with intermediate clustering of herds. Standard error for all values had an order of magnitude 10^−3^.

|  | Weak |  | Intermediate |  | Strong |  |
| --- | --- | --- | --- | --- | --- | --- |
|  | EBV | PBV | EBV | PBV | EBV | PBV |
| **Pedigree** |  |  |  |  |  |  |
| G | 0.54 | 0.43 | 0.65 | 0.40 | 0.70 | 0.37 |
| GH | 0.41 | 0.37 | 0.34 | 0.28 | 0.33 | 0.25 |
| GS | 0.17 | 0.17 | 0.17 | 0.18 | 0.18 | 0.18 |
| GHS | 0.16 | 0.16 | 0.17 | 0.18 | 0.18 | 0.18 |
| GHSC | 0.16 | 0.16 | 0.16 | 0.18 | 0.17 | 0.18 |
| **Genomic** |  |  |  |  |  |  |
| G | 0.39 | 0.39 | 0.32 | 0.30 | 0.30 | 0.26 |
| GH | 0.36 | 0.37 | 0.22 | 0.22 | 0.18 | 0.18 |
| GS | 0.15 | 0.15 | 0.13 | 0.15 | 0.13 | 0.15 |
| GHS | 0.14 | 0.15 | 0.12 | 0.15 | 0.12 | 0.14 |
| GHSC | 0.14 | 0.14 | 0.12 | 0.15 | 0.12 | 0.14 |

### Including environmental covariates

The environmental covariates did not improve the results substantially beyond the simple distance-driven spatial relationships between herds. This is shown for accuracy in Table 2 and CRPS in Table 3. Both the accuracy and CRPS were only marginally better for the GHSC model compared to the GHS model in some cases, and in the remaining cases they were comparable. Because of this we have focused on the sufficient models and excluded model GHSC in the remaining results. Some additional results with model GHSC are given in [Additional file 1].

### Separating genetic and spatial (environmental) effects

The models without spatial effects were not able to separate the genetic and spatial (environmental) effects. In Table 4 we present the correlation between the estimated breeding values and the true spatial effects by model and genetic connectedness. These results show that the models G and GH gave high correlation and suggest that the estimated breeding values captured parts of the spatial effects. The correlations from models GS and GHS were closer to zero, suggesting these models were better able to separate genetic and spatial effects. This, together with the correlation results in Table 2 and CRPS results in Table 3, suggests that the herd effect alone is not sufficient to account for all environmental effects in smallholder systems.

**Table 4:**
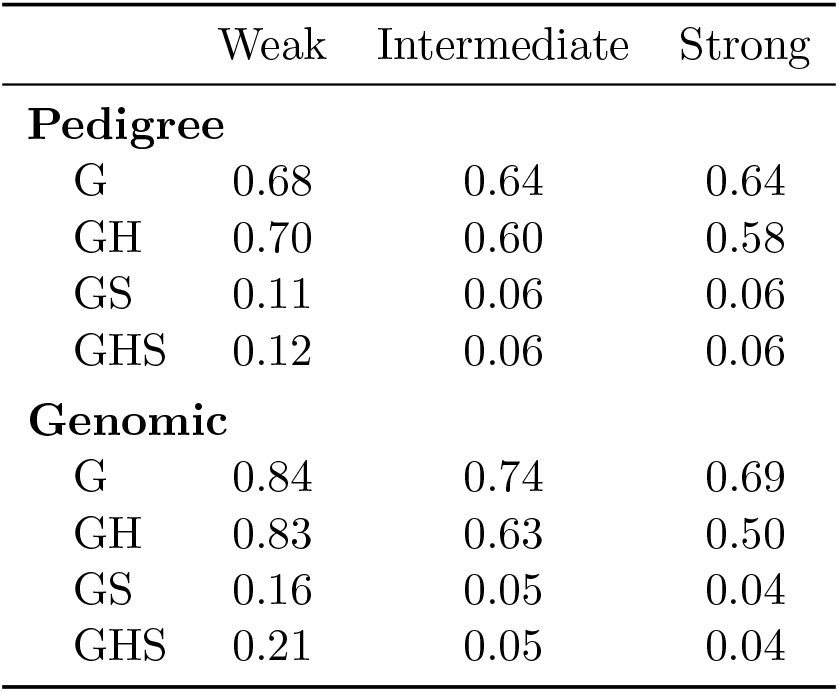
Average correlation between estimated breeding values and true spatial effect by genetic connectedness (weak, intermediate and strong) and model. Standard error for all values had an order of magnitude 10^−3^

### Comparing genetic connectedness scenarios and genetic models

The benefit of spatial modelling was largest when the spatial and genetic effects were hard to separate. In the [Additional file 1, Fig. 1] we show the relative improvement in accuracy and CRPS for model GH to model GHS by genetic connectedness. With both the genome and pedigree data the improvement was largest with weak genetic connectedness (about 50% to 80%), second with intermediate genetic connectedness (about 35% to 65%), and third with strong genetic connectedness (about 20% to 45%). These settings range between strongly confounded genetic and spatial effects to separable genetic and spatial effects. With weak genetic connectedness there was not much difference in improvement between models using genomic or pedigree data, whereas there was a tendency with intermediate and strong genetic connectedness that the improvement was largest with the pedigree data.

### Changing proportion of spatial variance

Spatial modelling in addition to the independent random herd effect, even when there were no spatial effects, did not decrease the accuracy. In Figure 2 we present the accuracy and CRPS for estimated breeding values when using genomic data under intermediate genetic connectedness. The *x*-axis goes from all phenotypic variance covered by herd effects to all covered by spatial effects. For models G and GH, the accuracy and CRPS worsened as the proportion of spatial variance increased, whereas for models GS and GHS the accuracy and CRPS improved. Overall, model GHS had the best accuracy and CRPS for all spatial variance proportions, and it was equal to model GH when there was no spatial variation and to model GS when there was no herd effect variation.

From the results so far we have seen that model GS had better accuracy and CRPS than model GH. However, this is not always the case. When most of the environmental variation was due to herd effects rather than spatial effects, model GH gave better estimates than model GS.

The same tendencies were seen for the predicted breeding values for both genomic and pedigree based models, and in other genetic connectedness scenarios, which can be seen in tables presented in [Additional file 1].

### Changing the herd clustering

When spatial and genetic effects were confounded the estimation accuracy improved when herds were weakly clustered rather than strongly clustered. When simulating the data we varied the distribution of herd locations, from strongly clustered within each village to less strongly clustered within each village. In Figure 3 we present the accuracy and CRPS for estimated breeding values using genomic data under weak genetic connectedness for the three clustering intensities. The figure shows that as herds were less clustered, the accuracy and CRPS improved across all models. We observed the same trend for predicted breeding values and using pedigree data, but not with intermediate and strong genetic connectedness, where the genetic and spatial effects were less confounded. Tables showing the accuracy and CRPS between true and inferred breeding values and correlation between inferred breeding values and true spatial effects for all levels of genetic connectedness and herd clustering are given in the [Additional file 1].

**Figure 3:**
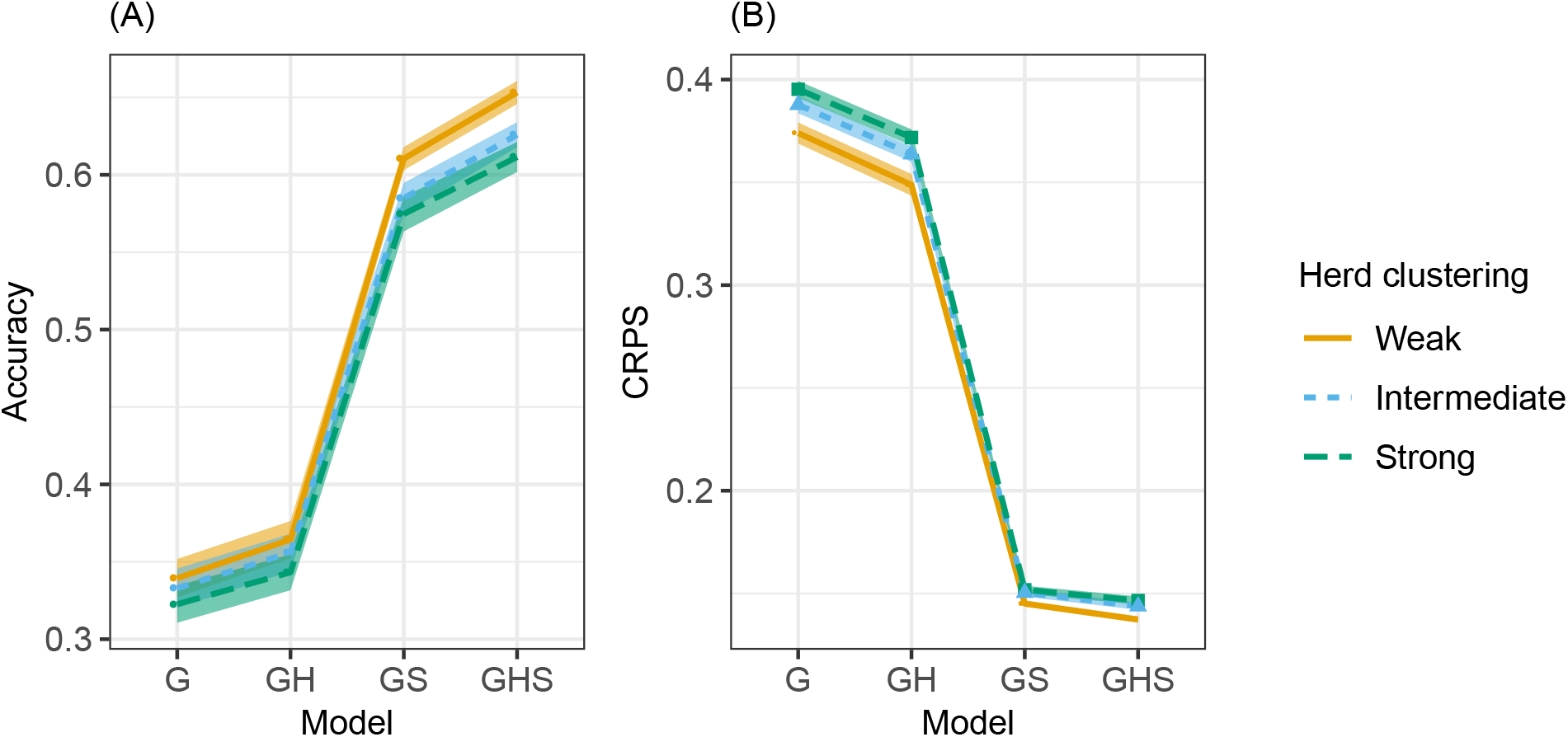
Average accuracy (A) and CRPS (smaller is better) (B) with 95% confidence intervals by model and herd clustering in scenario with weak genetic connectedness and using genomic model

### Real data

In this section we present the results from fitting the models to the subset of real cattle data. We present the posterior distributions of the hyper-parameters, the DIC, the estimated spatial field from model GHS, and compare the estimated breeding values from models GH and GHS. The corresponding results for the full data set are presented in the [Additional file 1]. Overall, the results showed that (1) models GH and GHS explained most of the variation in the data and had the best fit, (2) the data had a spatially dependent structure captured by models GS and GHS, and (3) the two models with the best fit, GH and GHS, separated the genetic and environmental effects differently.

### Explained variation and model fit

Models GH and GHS explained most of the variation in the data and had the best fit according to the DIC. In Figure 4 we show the posterior distributions for model hyperparameters. The figure has five panels showing the additive genetic variance 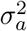, the residual variance 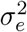, the herd effect variance 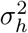, the spatial variance 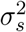, and the spatial range *ρ* in kilometers.

**Figure 4:**
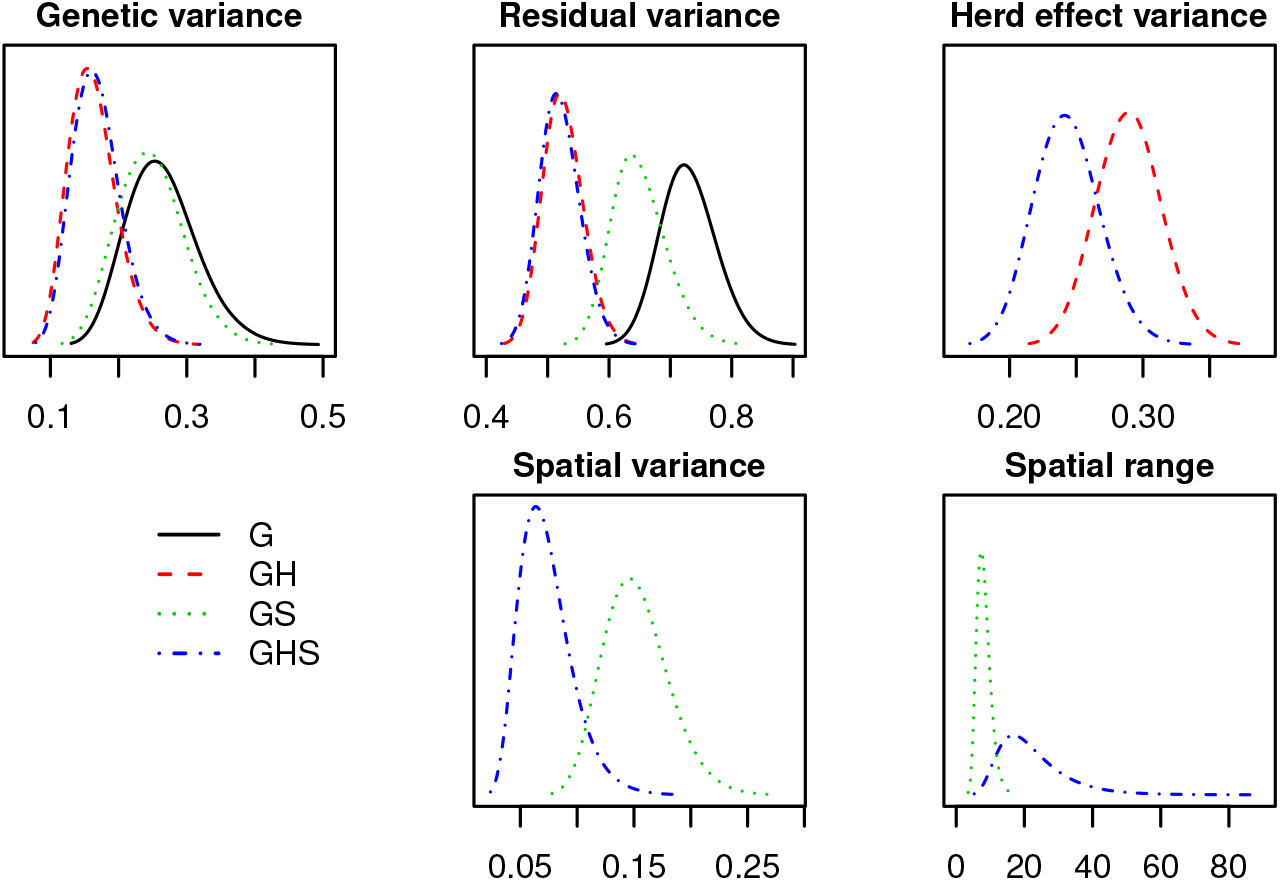
Posterior distributions of hyper-parameters from models G, GH, GS and GHS fitted to the real data

The posterior additive genetic variance was similar between models GH and GHS, higher in model GS, and even higher in model G. The same tendency was seen for the posterior residual variance. The posterior herd effect variance was lower in model GHS than model GH, which was reasonable since the herd effect in model GH captured the spatial component in the phenotype, which model GHS assigned to the spatial effect. The posterior spatial variance in model GS was higher than in model GHS since model GS captured herd effects. Finally, the posterior spatial range was lower in model GS than in model GHS, since model GS captured herd effects in the spatial effects which means shorter range of dependency between spatial locations. The mean posterior range from model GHS indicated that herds more than 22 km apart had close to independent (large scale) environments.

Since model G cannot explain variation due to herd or other environmental effects, it was reasonable to assume that some of the estimated genetic effect in G was actually due to confounding with other effects, which explains the high estimated additive genetic variance for this model. A similar reasoning could be used for model GS, which assigned variation due to herd effects, either to the genetic effects, the residual effects or the spatial effects. From Figure 4 it can seem that the variation from herd effects was distributed to all other effects, which explains why the estimated additive genetic variance and estimated residual variance was higher in model GS than models GH and GHS, and why the estimated spatial variance was higher than in model GHS. It seems that models GH and GHS distributed variation similarly except for the herd effect which is expected to be higher in GH than in GHS.

In Table 5 we show the DIC for the models. The table indicates that model GHS had the best fit, followed by model GH, then model GS and finally model G. These numbers are in line with the estimated hyper-parameters, where we saw that model GHS and GH could explain most of the variation in the phenotype. Although model GS has the potential to explain much variation as well, it is forced to assign herd effects either to genetic or spatial effects. We saw from the results with the simulated data that model GS had a worse model fit than model GH when most of the environmental variation was due to herd effects, which seems to be the case here considering the low posterior spatial variance. Finally, model G was not able to separate the genetic and environmental effects, which leads to a poor model fit. A rule of thumb, is that a complex model should be preferred over a less complex model if the DIC is reduced for more than ten units. When it comes to choosing between models GH and GHS, model GHS should be preferred, as its DIC was 36 units smaller.

**Table 5:**
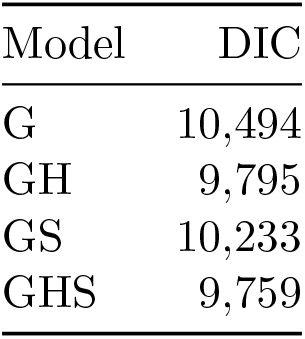
Deviance Information Criterion (DIC) by model fitted to the real data

| Model | DIC |
| --- | --- |
| G | 10,494 |
| GH | 9,795 |
| GS | 10,233 |
| GHS | 9,759 |

### The estimated spatial effects

The data had a spatially dependent structure captured by models GS and GHS, and the estimated spatial field from model GHS is shown in Figure 5. The figure shows both the estimated mean, in panel (A), and its uncertainty (posterior standard deviation) in panel (B). The axes show coordinates in the Transverse Mercator coordinate system in kilometres using datum WGS84.

**Figure 5:**
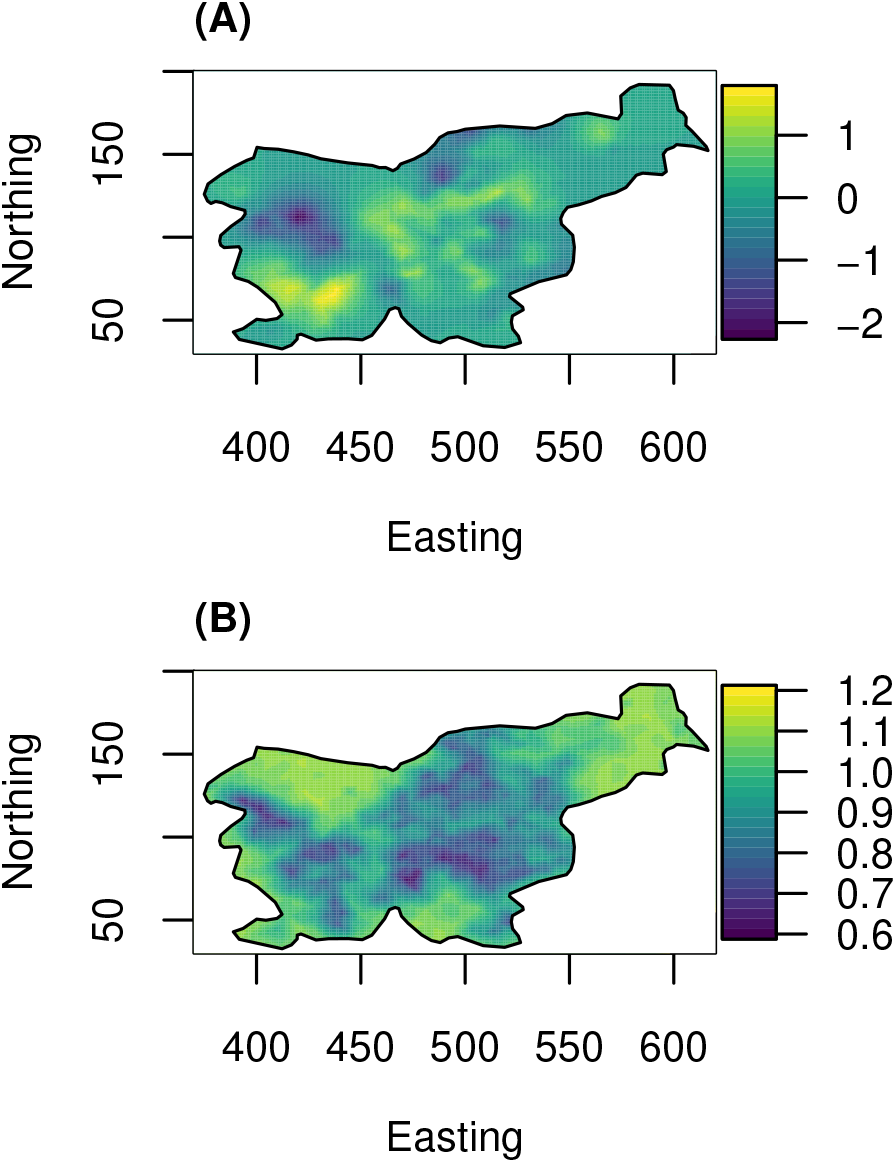
Posterior mean (A) and standard deviation (B) of the estimated spatial effect (in units of posterior spatial standard deviation) from model GHS fitted to the real data − the axis units are in km

In the western part of Slovenia model GHS suggest two environmental regions with mean different from zero, one with positive effect, and one with negative effect. In the central part of Slovenia, there are several smaller regions with either positive or negative environmental effect. In the north east there were not many observations, so there is only a small region of positive effect, and zero effects otherwise. These estimates are in line with the natural geographic conditions in Slovenia. The magnitude of these spatial effects range from ton −2.2 to 1.7 spatial standard deviations. The standard deviation was lowest where we had observations and was highest where there were no observations.

### Comparing breeding values from models GH and GHS

The two models with the best fit, models GH and GHS, separated the genetic and environmental effects differently. The DIC in Table 5 and the estimated hyper-parameters in Figure 4, indicated that models GH and GHS had the best model fit and a similar decomposition of the genetic and environmental effects. Furthermore, the estimated breeding values from models GH and GHS were highly correlated, with a correlation of about 0.995.

To evaluate how well the models separated the genetic and environmental effects, we computed the correlation of the estimated breeding values from models GH and GHS with the estimated (posterior mean) spatial effects from model GHS. For model GH this correlation was about 0.14, whereas for model GHS it was about 0.07. This suggests that there were some effects that were assigned as genetic in model GH, but assigned as spatial in model GHS.

In Figure 6 we present the difference in estimated breeding values between models GH and GHS as boxplots according to the estimated spatial effects from model GHS. This shows that the difference in estimated breeding values were correlated with the spatial effect from GHS. When the estimated spatial effects were negative, the estimated genetic effect from model GH was smaller than in model GHS, and when the estimated spatial effect was positive the genetic effect from model GH was larger than from model GHS. The magnitude of difference was in the range from −0.2 to 0.2 genetic standard deviations, which indicates substantial confounding. The figure also shows how many cows were used for each boxplot, which indicates that for many of the cows living in areas with intermediate spatial effects, the difference in estimated breeding values was not large.

**Figure 6:**
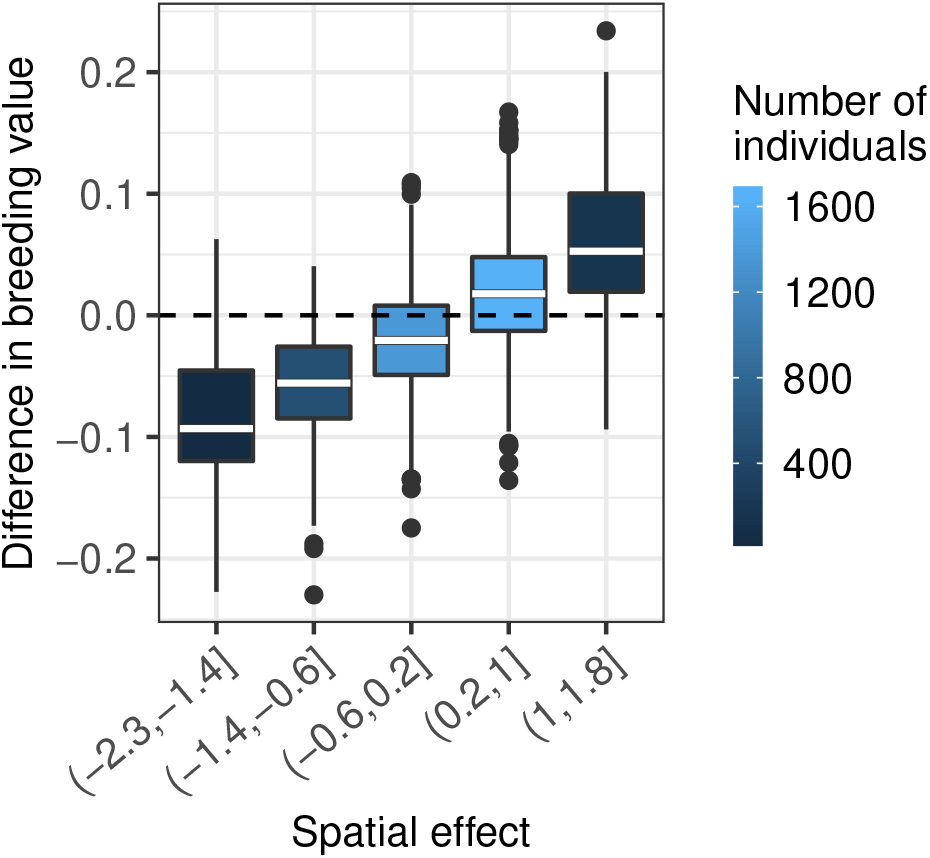
The difference in estimated breeding values (in units of posterior genetic standard deviation) between models GH and GHS by the estimated spatial effect (in units of posterior spatial standard deviation) from model GHS fitted to the real data

The correlation between the estimated breeding value differences and the estimated spatial effect from model GHS was about 0.62. This is in line with what we have seen from the simulation results and suggests that although the two models had highly correlated estimated breeding values, there were differences between the estimated breeding values due to model GH not separating the environmental and genetic effects as well as model GHS in the smallholders systems.

## Discussion

The results show that spatial modelling improves genetic evaluation in smallholder systems. In particular it increases the accuracy of genetic evaluation under weak genetic connectedness by establishing environmental connectedness and with this better separation of genetic and environmental effects. These observations highlight two broad points for discussion: (1) why spatial modelling improves genetic evaluation and (2) the limitations of this study and future possibilities.

### Why spatial modelling improves genetic evaluation

Spatial modelling improves genetic evaluation because it separates the environmental variation that is common for nearby herds from the phenotype. Since the spatial effects are estimated jointly for all herds and also jointly with other effects, such as management (herd) and genetic effects, it induces environmental connectedness and with this improves genetic connectedness. Animal breeders are very aware of the required data structures for accurate genetic evaluation (Foulley et al., 1990; Jorjani et al., 2001; Powell et al., 2019) and there are formal methods to asses genetic connectedness between contemporary groups (Foulley et al., 1992; Kennedy and Trus, 1993; Laloë, 1993; Laloë et al., 1996; Yu and Morota, 2019). Achieving sufficient genetic connectedness is particularly difficult when contemporary groups are small and there is limited genetic exchange between them.

A way to increase genetic connectedness is to use genomic data, though this was not sufficient in our case. Genomic data reveals genetic connectedness beyond pedigree data because animals likely share at least some alleles and this has been shown to increase accuracy of genetic evaluation (Powell et al., 2019; Yu et al., 2017, 2018). However, our targeted setting were smallholder herds, which are an extreme case of challenging data structure for genetic evaluation. Further, we varied genetic connectedness between villages. We found that across all genetic connectedness scenarios spatial modelling increased accuracy more than using genomic data instead of pedigree data. Further, with weakest genetic connectedness genomic data was not effective at all, while spatial modelling was. This is in a way not surprising because our herds were so small that we had strong confounding between genetic and environmental effects as well as weak genetic connectedness. So genomic data could not separate genetic and environmental effects and herds were too small to estimate their effect. In this case spatial modelling at least environmentally connected nearby herds and created effective contemporary groups. Accuracy was expectedly low in this extreme setting, though surprisingly not very low (see the next sub-section on possible reasons). These scenarios might seem too extreme, but they are a reflection of real situation in many countries around the world (e.g., Lawrence et al., 2015)

Spatial modelling has a long tradition and has been used before in animal breeding (e.g., Sæbø and Frigessi, 2004; Tiezzi et al., 2017). We have used it in the extreme scenario of small herds and for this reason choose to use the geostatistical approach to use all the available information. An alternative approach could be to cluster the small herds into village contemporary groups and potentially cluster these into region contemporary groups. In this case we would model village contemporary groups as an independent fixed or random effect and possibly model region contemporary groups as dependent random effect using the CAR model (Besag, 1974; Rue and Held, 2005). An issue with this approach is that we lose ability to model each individual herd and that administrative regions are sometimes a poor representation of geography. Given that the clustering approach has trade-offs, that there are efficient geostatistical models that adapt to data, and that efficient and easy to use implementations exist, we advise use of geostatistical models.

We advise routine use of spatial modelling in quantitative genetic models. Namely, collected data will always come from some area with likely variation in environmental effects. Our results show that spatial modelling is robust even when there is no spatial variation. The observed gains from this study will likely be smaller in cases with larger herds, but even in those cases spatial modelling can induce environmental connectedness and it can also provide estimates of spatial effects. These estimates could be used to target interventions or policies. Importantly, our analysis of simulated and real data collectively indicates that spatial modelling can remove spatial variation in estimated breeding values caused by environmental effects. Such modelling improvements will be very useful also beyond breeding populations; for example, in quantitative genetic analyses of human populations and wild populations. These populations also have similarly challenging data structure with rampant population structure (genetic disconnectedness) (Barton et al., 2019; Charmantier et al., 2014) and reports of biases in estimated genetic effects in line with geographic variation (Kerminen et al., 2019).

In line with the potential of spatial modelling to remove spatial variation we advise a geographically broad collection of data to train robust genomic models. Genomics is revolutionising breeding in developed and developing countries (VanRaden, 2008; Powell et al., 2019; Ojango et al., 2019). To deliver its full potential breeding organisations should ensure broad geographic coverage when collecting data. This will avoid bias towards a particular region. Spatial modelling can account for variation between and within regions, but it needs data from the regions to train optimal model parameters.

In relation to data collection guidance we were surprised to find that environmental covariates did not improve accuracy of genetic evaluation beyond simple distance-driven relationships between herds. To recall, we simulated the total spatial effect as a sum of eight spatial processes with a range of model parameters that made the processes quite different and we assumed that we can observe these with some noise. Our hypothesis was that modelling the observed environmental covariates would reveal the underlying spatial processes and increase accuracy in the same way that observed genomic data reveals the underlying genetic processes behind the pedigree expectations (VanRaden, 2008; Gorjanc et al., 2018). There are multiple possible explanations for this. Maybe we simulated too few spatial processes and the distance-based relationships was sufficient to model spatial variation. Maybe the noise in observations was too large or our data set was too small. Maybe the two-dimensional form of the space constrains the value of environmental covariates for increasing accuracy beyond the distance-based relationships. More studies are needed to address this question.

### The limitations of this study and future possibilities

There is a huge number of possible scenarios and parameter combinations that we could have tested. For example we assumed the absence of non-additive genetic effects, absence of genotype-by-environment interaction, absence of data errors, and considered only a single trait and breed. Further, the animals were initially distributed to herds by random, and the farms using artificial insemination were also chosen by random. Such simplifications are likely to yield higher accuracies and CRPS than what could be expected in real smallholder systems. However, the analysis of real data corroborates the main conclusions from the simulations. Future studies could for example consider non-random distribution of animals among herds as well as the use of artificial insemination and the best bulls. These non-random associations are realistic since well-resourced farmers are more likely to use artificial insemination and the best bulls (Schaeffer, 2018). With the real data that we applied in this paper, we tried to mimic a smallholder setting by using only a subset of the data set. However, note that this data has higher levels of artificial insemination than most smallholder systems.

Genotype-by-environment interactions have been modelled in several studies (Strandberg et al., 2009; Hayes et al., 2009; Tiezzi et al., 2017; Yao et al., 2017; Schultz and Weigel, 2019) and such interactions are likely substantial in smallholder systems, in particular when native and exotic breeds are used (Ojango et al., 2019). We ignored these interactions in our study. Of particular notice to these interactions and in relation to our work is the study of Tiezzi et al. (2017). They used geographical location and weather data in addition to herd summaries to describe environmental conditions in genetic evaluations, with and without genotype-by-environment interaction and concluded that the farming environment explained variation in the data, as well as the genotype-by-environment component. Further work is needed to embrace the rich set of models and tools from the spatial statistics community to address genotype-by-environment interactions (Heaton et al., 2019; van Niekerk et al., 2019).

Yet another important source of phenotypic variation that we ignored are heterogeneous variances, which are likely substantial in smallholder systems. There are multiple models and methods used by breeders and geneticists to account for such variation (e.g., Wiggans and VanRaden, 1991; Visscher and Hill, 1992; Meuwissen et al., 1996). We note that there is also a rich spatial literature on models that can deal with non-stationarity in dependency and variance (e.g., Sampson and Guttorp, 1992; Fuentes, 2001; Higdon, 2002; Lindgren et al., 2011; Ingebrigtsen et al., 2014), which for example could enable modelling directional dependence based on local anisotropy (e.g., Fuglstad et al., 2015a). Using and benefiting from non-stationary models can challenging due to compuational costs and the amount of data needed to fit the models (Fuglstad et al., 2015b). But this will be increasingly possible and desired as the data sets increase in size with the progression of digital revolution in agriculture and more computationally efficient methods become available.

Breeding programmes interested in spatial modelling will have to invest in software modification. This is not a limitation of this study, but interested breeding programmes would either have to use the R-INLA package (Rue et al., 2009) or implement extension of their existing software. While the R-INLA package is a mature project it does not support all the models that animal breeders use, most notably the multi-trait models. However, it handles a rich set of likelihoods (Gaussian, Poisson, Bernoulli, Weibull, etc.), link functions, independent or correlated random effects (time-series, regions, points, generic such as pedigree, etc.) and priors. It uses the same key underlying linear algebra routines as standard genetic evaluation software (Takahashi, 1973; Rue and Held, 2005; De Coninck et al., 2016; Verbosio et al., 2017) and enables both full Bayesian analysis with fast and very accurate approximate algorithm (Rue and Martino, 2007) or even faster empirical Bayesian analysis. We have used it extensively for standard quantitative genetic studies (Holand et al., 2013; Larsen et al., 2014; Muff et al., 2019), accounting for selection (Steinsland et al., 2014), spatial modelling of plant and tree trials (Selle et al., 2019) and for modelling of phenotypes on phylogeny (Selle et al., 2020). While the R-INLA package is fast for models with sparse structure (time-series, spatial regions or points and pedigree) it does not fare well for genomic models that have dense structure VanRaden (2008); Gorjanc et al. (2018). However, use of recently proposed approximate genomic models (Misztal et al., 2014; Misztal, 2016) and sparse-dense libraries would help (Masuda et al., 2014, 2015). A simple alternative for spatial modelling with stahdard software would be to brute force setup and inversion of the spatial covariance matrix (using Matérn or Gaussian covariance). This would suffice for a few thousand well dispersed herds, but might lead to numeric issues with nearby herds (near matrix singularity) or much larger numbers of herds that will soon become reality with digital revolution of agriculture.

Further, since INLA does a full Bayesian analysis, the user has to set prior distributions for all model parameters. This is not always straightforward, but setting a prior based on the knowledge about the process is likely to improve inference substantially. There is a number of ways to set mildly informative priors. We used penalised complexity priors (Simpson et al., 2017) since these avoid over-fitting and can accommodate prior knowledge about the relative importance of the different effects in the models (Fuglstad et al., 2018; Hem et al., 2020).

## Conclusions

This study demonstrates that spatial modelling can improve genetic evaluation in smallholder systems by inducing environmental connectedness and with this better separation of genetic and environmental effects beyond an independent herd effect. We have demonstrated this with simulated data with different levels of genetic connectedness, proportions of spatial to management (herd) variation, herd clustering and pedigree or genomic modelling. Spatial modelling in addition to a herd effect did not perform worse than a model with only a herd effect even when there was no spatial effect. We expected that environmental covariates would improve spatial modelling following the analogy of genetic modelling with observed genomic vs. expected pedigree data, but this was not the case, possibly due to a too simple simulation and inherent limitation that space is twodimensional. We have also shown with real data that spatial modelling separated genetic and environmental effects differently and that the difference correlated with the spatial effect. The estimated spatial effects can also be used to guide management and polices. Based on all of these results we suggest routine spatial modelling in genetic evaluations, particularly for smallholder systems. Spatial modelling could also have major impact in studies of human and wild populations.

## Additional Files

**Additional file 1 – Additional results**

**Additional file 2 – Simulation code available from** https://doi.org/10.6084/m9.figshare.12403898

