## Additional file 1 for "Spatial modelling improves genetic evaluation in smallholder breeding programs"

June 1, 2020

### Simulated data

#### Comparing genetic connectedness and genetic models

In Figure 1 we plot the relative improvement in average accuracy and CRPS between true breeding values (TBV) and estimated breeding values (EBV) or predicted breeding values (PBV) when using models GH or GHS, for the different levels of genetic connectedness.

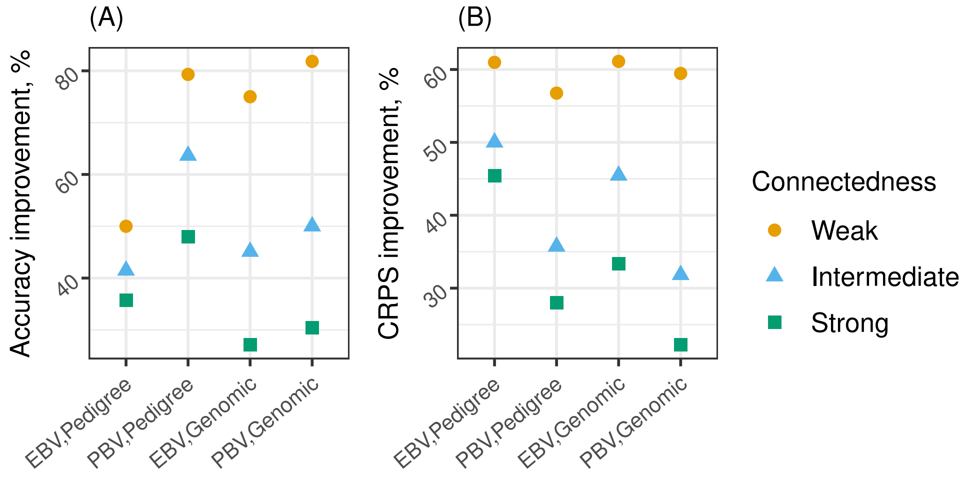


Figure 1: Percentage improvement in EBV accuracy (A) and CRPS (B) between models GH and GHS by genetic connectedness and genetic model

#### Changing proportion of spatial variance

Here we show the average accuracy and CRPS between TBV and EBV or PBV for all levels of genetic connectedness, using both pedigree and genomic data, when varying the proportion of spatial variance relative to the sum of spatial variance and herd effect variance. The herd locations were simulated from a bi-variate normal distribution with mean equal to the village locations, and variance $3.5 \cdot 10^{-4} \boldsymbol{I}_{2\times 2}$ (intermediate herd clustering).

Table 1 and Table 2 respectively show accuracy and CRPS for weak genetic connectedness. Table 3 and Table 4 respectively show accuracy and CRPS for intermediate genetic connectedness. Table 5 and Table 6 respectively show accuracy and CRPS for strong genetic connectedness.

Table 1: Average accuracy for EBV and PBV with weak genetic connectedness, using pedigree or genomic data, with varying proportion of spatial variance. The standard error for some values had order of magnitude $10^{-2}$, and most had $10^{-3}$

| | EBV | | | | | | PBV | | | | | |
|---|---|---|---|---|---|---|---|---|---|---|---|---|
| $\sigma_s^2/(\sigma_s^2+\sigma_h^2)$ | 0 | 0.2 | 0.4 | 0.6 | 0.8 | 1 | 0 | 0.2 | 0.4 | 0.6 | 0.8 | 1 |
| **Pedigree** | | | | | | | | | | | | |
| G | 0.53 | 0.40 | 0.35 | 0.34 | 0.32 | 0.31 | 0.54 | 0.38 | 0.32 | 0.29 | 0.26 | 0.23 |
| GH | 0.57 | 0.46 | 0.39 | 0.37 | 0.34 | 0.32 | 0.57 | 0.42 | 0.34 | 0.31 | 0.27 | 0.24 |
| GS | 0.51 | 0.48 | 0.49 | 0.52 | 0.56 | 0.60 | 0.51 | 0.46 | 0.47 | 0.50 | 0.53 | 0.58 |
| GHS | 0.57 | 0.53 | 0.53 | 0.54 | 0.56 | 0.60 | 0.56 | 0.51 | 0.51 | 0.52 | 0.53 | 0.58 |
| **Genomic** | | | | | | | | | | | | |
| G | 0.54 | 0.40 | 0.36 | 0.34 | 0.31 | 0.32 | 0.55 | 0.38 | 0.34 | 0.31 | 0.27 | 0.30 |
| GH | 0.64 | 0.47 | 0.40 | 0.36 | 0.33 | 0.33 | 0.62 | 0.43 | 0.37 | 0.33 | 0.28 | 0.30 |
| GS | 0.53 | 0.51 | 0.54 | 0.57 | 0.63 | 0.70 | 0.53 | 0.50 | 0.53 | 0.55 | 0.60 | 0.67 |
| GHS | 0.64 | 0.60 | 0.61 | 0.62 | 0.64 | 0.70 | 0.62 | 0.58 | 0.58 | 0.58 | 0.61 | 0.67 |

Table 2: Average CRPS for EBV and PBV with weak genetic connectedness, using pedigree or genomic data, with varying proportion of spatial variance. The standard error for all values had order of magnitude $10^{-3}$

| | EBV | | | | | | PBV | | | | | |
|---|---|---|---|---|---|---|---|---|---|---|---|---|
| $\sigma_s^2/(\sigma_s^2+\sigma_h^2)$ | 0 | 0.2 | 0.4 | 0.6 | 0.8 | 1 | 0 | 0.2 | 0.4 | 0.6 | 0.8 | 1 |
| **Pedigree** | | | | | | | | | | | | |
| G | 0.164 | 0.271 | 0.395 | 0.552 | 0.688 | 0.730 | 0.164 | 0.255 | 0.347 | 0.429 | 0.489 | 0.542 |
| GH | 0.155 | 0.237 | 0.332 | 0.406 | 0.492 | 0.618 | 0.155 | 0.213 | 0.289 | 0.359 | 0.431 | 0.519 |
| GS | 0.161 | 0.170 | 0.171 | 0.170 | 0.168 | 0.165 | 0.162 | 0.155 | 0.155 | 0.152 | 0.149 | 0.142 |
| GHS | 0.155 | 0.164 | 0.167 | 0.166 | 0.166 | 0.164 | 0.155 | 0.146 | 0.148 | 0.147 | 0.147 | 0.142 |
| **Genomic** | | | | | | | | | | | | |
| G | 0.158 | 0.242 | 0.317 | 0.385 | 0.452 | 0.514 | 0.158 | 0.237 | 0.313 | 0.384 | 0.450 | 0.524 |
| GH | 0.139 | 0.214 | 0.287 | 0.359 | 0.436 | 0.508 | 0.142 | 0.209 | 0.284 | 0.357 | 0.433 | 0.517 |
| GS | 0.154 | 0.160 | 0.157 | 0.153 | 0.144 | 0.129 | 0.155 | 0.141 | 0.142 | 0.138 | 0.131 | 0.116 |
| GHS | 0.139 | 0.148 | 0.147 | 0.146 | 0.140 | 0.129 | 0.142 | 0.134 | 0.135 | 0.133 | 0.127 | 0.116 |

Table 3: Average accuracy for EBV and PBV with intermediate genetic connectedness, using pedigree or genomic data, with varying proportion of spatial variance. The standard error for some values had order of magnitude $10^{-2}$, and most had $10^{-3}$

| | EBV | | | | | | PBV | | | | | |
|---|---|---|---|---|---|---|---|---|---|---|---|---|
| $\sigma_s^2/(\sigma_s^2+\sigma_h^2)$ | 0 | 0.2 | 0.4 | 0.6 | 0.8 | 1 | 0 | 0.2 | 0.4 | 0.6 | 0.8 | 1 |
| **Pedigree** | | | | | | | | | | | | |
| G | 0.51 | 0.40 | 0.35 | 0.33 | 0.32 | 0.32 | 0.29 | 0.22 | 0.18 | 0.17 | 0.17 | 0.16 |
| GH | 0.61 | 0.51 | 0.44 | 0.39 | 0.36 | 0.33 | 0.36 | 0.29 | 0.24 | 0.21 | 0.18 | 0.16 |
| GS | 0.53 | 0.53 | 0.55 | 0.59 | 0.61 | 0.65 | 0.31 | 0.32 | 0.33 | 0.35 | 0.37 | 0.41 |
| GHS | 0.61 | 0.60 | 0.60 | 0.62 | 0.62 | 0.65 | 0.37 | 0.37 | 0.37 | 0.38 | 0.38 | 0.41 |
| **Genomic** | | | | | | | | | | | | |
| G | 0.61 | 0.51 | 0.42 | 0.39 | 0.37 | 0.35 | 0.46 | 0.40 | 0.33 | 0.26 | 0.26 | 0.26 |
| GH | 0.72 | 0.66 | 0.57 | 0.51 | 0.45 | 0.39 | 0.56 | 0.52 | 0.44 | 0.36 | 0.33 | 0.28 |
| GS | 0.63 | 0.65 | 0.67 | 0.70 | 0.73 | 0.77 | 0.47 | 0.52 | 0.54 | 0.54 | 0.57 | 0.60 |
| GHS | 0.72 | 0.72 | 0.72 | 0.73 | 0.74 | 0.77 | 0.56 | 0.58 | 0.58 | 0.57 | 0.58 | 0.60 |

Table 4: Average CRPS for EBV and PBV with intermediate genetic connectedness, using pedigree or genomic data, with varying proportion of spatial variance. The standard error for all values had order of magnitude $10^{-3}$

| | EBV | | | | | | PBV | | | | | |
|---|---|---|---|---|---|---|---|---|---|---|---|---|
| $\sigma_s^2/(\sigma_s^2+\sigma_h^2)$ | 0 | 0.2 | 0.4 | 0.6 | 0.8 | 1 | 0 | 0.2 | 0.4 | 0.6 | 0.8 | 1 |
| **Pedigree** | | | | | | | | | | | | |
| G | 0.165 | 0.268 | 0.396 | 0.536 | 0.685 | 0.709 | 0.181 | 0.233 | 0.306 | 0.364 | 0.411 | 0.431 |
| GH | 0.153 | 0.211 | 0.296 | 0.370 | 0.455 | 0.578 | 0.174 | 0.189 | 0.239 | 0.284 | 0.335 | 0.398 |
| GS | 0.160 | 0.167 | 0.164 | 0.159 | 0.159 | 0.153 | 0.178 | 0.160 | 0.158 | 0.156 | 0.154 | 0.150 |
| GHS | 0.153 | 0.161 | 0.159 | 0.157 | 0.158 | 0.153 | 0.173 | 0.156 | 0.154 | 0.154 | 0.153 | 0.150 |
| **Genomic** | | | | | | | | | | | | |
| G | 0.145 | 0.201 | 0.262 | 0.319 | 0.372 | 0.428 | 0.162 | 0.187 | 0.235 | 0.285 | 0.322 | 0.364 |
| GH | 0.126 | 0.144 | 0.176 | 0.218 | 0.276 | 0.358 | 0.148 | 0.141 | 0.165 | 0.203 | 0.244 | 0.311 |
| GS | 0.141 | 0.139 | 0.135 | 0.131 | 0.124 | 0.114 | 0.159 | 0.134 | 0.129 | 0.130 | 0.125 | 0.120 |
| GHS | 0.126 | 0.125 | 0.125 | 0.124 | 0.121 | 0.114 | 0.148 | 0.125 | 0.123 | 0.125 | 0.122 | 0.120 |

Table 5: Average accuracy for EBV and PBV with strong genetic connectedness, using pedigree or genomic data, with varying proportion of spatial variance. The standard error for some values had order of magnitude $10^{-2}$, and most had $10^{-3}$

| | EBV | | | | | | PBV | | | | | |
|---|---|---|---|---|---|---|---|---|---|---|---|---|
| $\sigma_s^2/(\sigma_s^2+\sigma_h^2)$ | 0 | 0.2 | 0.4 | 0.6 | 0.8 | 1 | 0 | 0.2 | 0.4 | 0.6 | 0.8 | 1 |
| **Pedigree** | | | | | | | | | | | | |
| G | 0.49 | 0.38 | 0.32 | 0.32 | 0.31 | 0.32 | 0.30 | 0.24 | 0.20 | 0.19 | 0.18 | 0.17 |
| GH | 0.57 | 0.50 | 0.44 | 0.41 | 0.38 | 0.35 | 0.36 | 0.32 | 0.27 | 0.25 | 0.22 | 0.19 |
| GS | 0.51 | 0.52 | 0.53 | 0.55 | 0.57 | 0.61 | 0.31 | 0.33 | 0.34 | 0.35 | 0.36 | 0.39 |
| GHS | 0.57 | 0.57 | 0.57 | 0.57 | 0.58 | 0.61 | 0.36 | 0.37 | 0.37 | 0.37 | 0.37 | 0.39 |
| **Genomic** | | | | | | | | | | | | |
| G | 0.65 | 0.54 | 0.49 | 0.44 | 0.40 | 0.38 | 0.52 | 0.41 | 0.38 | 0.35 | 0.27 | 0.29 |
| GH | 0.74 | 0.69 | 0.66 | 0.60 | 0.55 | 0.49 | 0.60 | 0.54 | 0.52 | 0.48 | 0.39 | 0.38 |
| GS | 0.67 | 0.67 | 0.70 | 0.72 | 0.75 | 0.79 | 0.53 | 0.53 | 0.56 | 0.58 | 0.59 | 0.63 |
| GHS | 0.74 | 0.74 | 0.75 | 0.75 | 0.76 | 0.79 | 0.60 | 0.59 | 0.61 | 0.60 | 0.60 | 0.63 |

Table 6: Average CRPS for EBV and PBV with strong genetic connectedness, using pedigree or genomic data, with varying proportion of spatial variance. The standard error for all values had order of data $10^{-3}$

| | EBV | | | | | | PBV | | | | | |
|---|---|---|---|---|---|---|---|---|---|---|---|---|
| $\sigma_s^2/(\sigma_s^2+\sigma_h^2)$ | 0 | 0.2 | 0.4 | 0.6 | 0.8 | 1 | 0 | 0.2 | 0.4 | 0.6 | 0.8 | 1 |
| **Pedigree** | | | | | | | | | | | | |
| G | 0.180 | 0.329 | 0.555 | 0.701 | 0.720 | 0.731 | 0.187 | 0.247 | 0.331 | 0.363 | 0.375 | 0.389 |
| GH | 0.171 | 0.234 | 0.288 | 0.336 | 0.383 | 0.499 | 0.179 | 0.189 | 0.217 | 0.239 | 0.267 | 0.324 |
| GS | 0.174 | 0.179 | 0.178 | 0.180 | 0.182 | 0.180 | 0.184 | 0.165 | 0.164 | 0.165 | 0.164 | 0.159 |
| GHS | 0.170 | 0.178 | 0.175 | 0.177 | 0.181 | 0.179 | 0.179 | 0.162 | 0.160 | 0.161 | 0.162 | 0.159 |
| **Genomic** | | | | | | | | | | | | |
| G | 0.139 | 0.188 | 0.237 | 0.289 | 0.337 | 0.394 | 0.155 | 0.173 | 0.211 | 0.243 | 0.289 | 0.313 |
| GH | 0.122 | 0.134 | 0.149 | 0.176 | 0.205 | 0.264 | 0.143 | 0.132 | 0.142 | 0.160 | 0.188 | 0.221 |
| GS | 0.136 | 0.135 | 0.130 | 0.126 | 0.120 | 0.111 | 0.154 | 0.130 | 0.127 | 0.125 | 0.122 | 0.115 |
| GHS | 0.122 | 0.122 | 0.120 | 0.120 | 0.117 | 0.111 | 0.143 | 0.122 | 0.120 | 0.122 | 0.120 | 0.115 |

### Changing the herd clustering

Table 7 and Table 8 respectively show average accuracy and CRPS between TBV and EBV/PBV for all levels of genetic connectedness, using both pedigree and genomic data, when the herd locations were simulated from a bi-variate normal distribution with mean equal to the village locations, and variance $1 \cdot 10^{-4} \boldsymbol{I}_{2\times 2}$ (strong herd clustering). Table 9 and Table 10 respectively show average accuracy and CRPS between TBV and EBV/PBV for all levels of genetic connectedness, using both pedigree and genomic data, when the herd locations were simulated from a bi-variate normal distribution with mean equal to the village locations, and variance $9 \cdot 10^{-4} \boldsymbol{I}_{2\times 2}$ (weak herd clustering).

Table 7: Average accuracy for the different levels of genetic connectedness for EBV and PBV, using pedigree or genomic data, and the herd locations simulated using variance $1 \cdot 10^{-4} \boldsymbol{I}_{2\times 2}$ (strong herd clustering). The standard error for some values had order of magnitude $10^{-2}$, and most had $10^{-3}$

| | Weak | | Intermediate | | Strong | |
|---|---|---|---|---|---|---|
| | EBV | PBV | EBV | PBV | EBV | PBV |
| **Pedigree** | | | | | | |
| G | 0.32 | 0.27 | 0.32 | 0.18 | 0.32 | 0.19 |
| GH | 0.35 | 0.29 | 0.41 | 0.22 | 0.41 | 0.25 |
| GS | 0.51 | 0.48 | 0.56 | 0.34 | 0.55 | 0.35 |
| GHS | 0.53 | 0.50 | 0.58 | 0.36 | 0.57 | 0.37 |
| GHSC | 0.56 | 0.54 | 0.59 | 0.37 | 0.58 | 0.38 |
| **Genomic** | | | | | | |
| G | 0.32 | 0.30 | 0.40 | 0.29 | 0.42 | 0.32 |
| GH | 0.34 | 0.32 | 0.51 | 0.38 | 0.58 | 0.46 |
| GS | 0.57 | 0.55 | 0.70 | 0.54 | 0.72 | 0.57 |
| GHS | 0.61 | 0.58 | 0.73 | 0.58 | 0.75 | 0.60 |
| GHSC | 0.63 | 0.60 | 0.74 | 0.59 | 0.75 | 0.61 |

Table 8: Average CRPS for different genetic connectedness for EBV and PBV, using pedigree or genomic data, and the herd locations simulated using variance $1 \cdot 10^{-4} \boldsymbol{I}_{2\times 2}$ (strong herd clustering). The standard error for all values had order of magnitude $10^{-3}$

| | Weak | | Intermediate | | Strong | |
|---|---|---|---|---|---|---|
| | EBV | PBV | EBV | PBV | EBV | PBV |
| **Pedigree** | | | | | | |
| G | 0.559 | 0.438 | 0.667 | 0.406 | 0.706 | 0.371 |
| GH | 0.419 | 0.374 | 0.343 | 0.281 | 0.335 | 0.252 |
| GS | 0.168 | 0.168 | 0.166 | 0.180 | 0.180 | 0.183 |
| GHS | 0.165 | 0.164 | 0.166 | 0.178 | 0.179 | 0.181 |
| GHSC | 0.160 | 0.159 | 0.163 | 0.176 | 0.176 | 0.179 |
| **Genomic** | | | | | | |
| G | 0.395 | 0.402 | 0.325 | 0.302 | 0.299 | 0.264 |
| GH | 0.372 | 0.378 | 0.225 | 0.222 | 0.180 | 0.181 |
| GS | 0.152 | 0.156 | 0.130 | 0.151 | 0.126 | 0.146 |
| GHS | 0.147 | 0.151 | 0.124 | 0.146 | 0.120 | 0.142 |
| GHSC | 0.143 | 0.147 | 0.123 | 0.145 | 0.120 | 0.142 |

Table 9: Average accuracy for the different levels of genetic connectedness for EBV and PBV, using pedigree or genomic data, and the herd locations simulated using variance $9 \cdot 10^{-4} \boldsymbol{I}_{2\times 2}$ (weak herd clustering). The standard error for some values had order of magnitude $10^{-2}$, and most had $10^{-3}$

| | Weak | | Intermediate | | Strong | |
|---|---|---|---|---|---|---|
| | EBV | PBV | EBV | PBV | EBV | PBV |
| **Pedigree** | | | | | | |
| G | 0.33 | 0.29 | 0.32 | 0.17 | 0.32 | 0.19 |
| GH | 0.37 | 0.31 | 0.41 | 0.22 | 0.42 | 0.25 |
| GS | 0.55 | 0.54 | 0.56 | 0.34 | 0.55 | 0.35 |
| GHS | 0.56 | 0.55 | 0.59 | 0.37 | 0.57 | 0.37 |
| GHSC | 0.58 | 0.57 | 0.59 | 0.37 | 0.58 | 0.37 |
| **Genomic** | | | | | | |
| G | 0.34 | 0.31 | 0.40 | 0.27 | 0.44 | 0.33 |
| GH | 0.36 | 0.33 | 0.53 | 0.38 | 0.61 | 0.47 |
| GS | 0.61 | 0.60 | 0.70 | 0.54 | 0.72 | 0.57 |
| GHS | 0.65 | 0.63 | 0.74 | 0.57 | 0.75 | 0.59 |
| GHSC | 0.67 | 0.65 | 0.74 | 0.58 | 0.75 | 0.60 |

Table 10: Average CRPS for the different levels of genetic connectedness for EBV and PBV, using pedigree or genomic data, and the herd locations simulated using variance $9 \cdot 10^{-4} \boldsymbol{I}_{2\times 2}$ (weak herd clustering). The standard error for all values had order of magnitude $10^{-3}$

| | Weak | | Intermediate | | Strong | |
|---|---|---|---|---|---|---|
| | EBV | PBV | EBV | PBV | EBV | PBV |
| **Pedigree** | | | | | | |
| G | 0.500 | 0.410 | 0.615 | 0.392 | 0.688 | 0.370 |
| GH | 0.393 | 0.348 | 0.326 | 0.270 | 0.325 | 0.248 |
| GS | 0.164 | 0.163 | 0.165 | 0.179 | 0.180 | 0.184 |
| GHS | 0.160 | 0.158 | 0.163 | 0.176 | 0.178 | 0.181 |
| GHSC | 0.156 | 0.155 | 0.161 | 0.175 | 0.177 | 0.180 |
| **Genomic** | | | | | | |
| G | 0.374 | 0.387 | 0.308 | 0.289 | 0.282 | 0.253 |
| GH | 0.349 | 0.362 | 0.209 | 0.211 | 0.169 | 0.174 |
| GS | 0.145 | 0.147 | 0.130 | 0.152 | 0.126 | 0.147 |
| GHS | 0.137 | 0.141 | 0.123 | 0.146 | 0.119 | 0.143 |
| GHSC | 0.135 | 0.138 | 0.122 | 0.146 | 0.119 | 0.143 |

### Correlation between true spatial effect and EBV with changing herd clustering

Table 11 shows the average correlation between the EBV and the true spatial effect for all levels of genetic connectedness, using pedigree or genomic data, when the herd locations were simulated from a bi-variate normal distribution with mean equal to the village locations, and variance $1 \cdot 10^{-4}\boldsymbol{I}_{2\times2}$ (strong herd clustering).

Table 12 shows the average correlation between the EBV and the true spatial effect for all levels of genetic connectedness, using pedigree or genomic data, when the herd locations were simulated from a bi-variate normal distribution with mean equal to the village locations, and variance $3.5 \cdot 10^{-4}\boldsymbol{I}_{2\times2}$ (intermediate herd clustering). This is an extended table from the main paper where model GHSC is included.

Table 13 shows the average correlation between the EBV and the true spatial effect for all levels of genetic connectedness, using pedigree or genomic data, when the herd locations were simulated from a bi-variate normal distribution with mean equal to the village locations, and variance $9 \cdot 10^{-4}\boldsymbol{I}_{2\times2}$ (weak herd clustering).

Table 11: Average correlation between EBV and true spatial effect for all levels of genetic connectedness, using pedigree or genomic data, and the herd locations simulated using variance $1 \cdot 10^{-4}\boldsymbol{I}_{2\times2}$ (strong herd clustering). The standard error for all values had order of magnitude $10^{-3}$

| Connectedness | Weak | Intermediate | Strong |
|---|---|---|---|
| **Pedigree** | | | |
| G | 0.87 | 0.76 | 0.70 |
| GH | 0.86 | 0.65 | 0.51 |
| GS | 0.23 | 0.06 | 0.03 |
| GHS | 0.27 | 0.07 | 0.03 |
| GHSC | 0.21 | 0.06 | 0.03 |
| **Genomic** | | | |
| G | 0.67 | 0.64 | 0.63 |
| GH | 0.71 | 0.62 | 0.60 |
| GS | 0.12 | 0.07 | 0.06 |
| GHS | 0.13 | 0.06 | 0.06 |
| GHSC | 0.10 | 0.05 | 0.05 |

Table 12: Average correlation between EBV and true spatial effect for all levels of genetic connectedness, using pedigree or genomic data, and the herd locations simulated using variance $3.5 \cdot 10^{-4}\boldsymbol{I}_{2\times2}$ (intermediate herd clustering). The standard error for all values had order of magnitude $10^{-3}$

| Connectedness | Weak | Intermediate | Strong |
|---|---|---|---|
| **Pedigree** | | | |
| G | 0.68 | 0.64 | 0.64 |
| GH | 0.70 | 0.60 | 0.58 |
| GS | 0.11 | 0.06 | 0.06 |
| GHS | 0.12 | 0.06 | 0.06 |
| GHSC | 0.10 | 0.06 | 0.06 |
| **Genomic** | | | |
| G | 0.84 | 0.74 | 0.69 |
| GH | 0.83 | 0.63 | 0.50 |
| GS | 0.16 | 0.05 | 0.04 |
| GHS | 0.21 | 0.05 | 0.04 |
| GHSC | 0.18 | 0.05 | 0.03 |

Table 13: Average correlation between EBV and true spatial effect for all levels of genetic connectedness, using pedigree or genomic data, and the herd locations simulated using variance $9 \cdot 10^{-4}\boldsymbol{I}_{2\times 2}$ (weak herd clustering). The standard error for all values had order of magnitude $10^{-3}$

| Connectedness | Weak | Intermediate | Strong |
|---|---|---|---|
| **Pedigree** | | | |
| G | 0.83 | 0.72 | 0.67 |
| GH | 0.81 | 0.59 | 0.47 |
| GS | 0.12 | 0.04 | 0.03 |
| GHS | 0.15 | 0.04 | 0.03 |
| GHSC | 0.14 | 0.04 | 0.02 |
| **Genomic** | | | |
| G | 0.69 | 0.65 | 0.63 |
| GH | 0.67 | 0.57 | 0.56 |
| GS | 0.09 | 0.05 | 0.05 |
| GHS | 0.09 | 0.05 | 0.05 |
| GHSC | 0.08 | 0.04 | 0.04 |

### Real data

The herd locations of the 1,838 different herds in Slovenia are shown in Figure 2. The axes show the coordinates in kilometres in the Transverse Mercator coordinate system using datum WGS84.

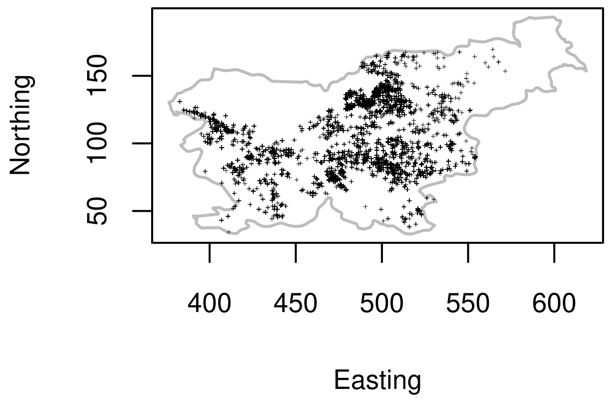


Figure 2: The location of the herds in the BSC data shown with black points, and the border of Slovenia in grey. The axis units are in km

For the models G, GH, GS and GHS applied to the full real data, we present the posterior hyper-parameters in Figure 3, the DIC in Table 14, the posterior mean and standard deviation of the estimated spatial effects from model GHS in Figure 4, and the difference in EBV between models GH and GHS plotted against the mean posterior spatial effect from model GHS in Figure 5.

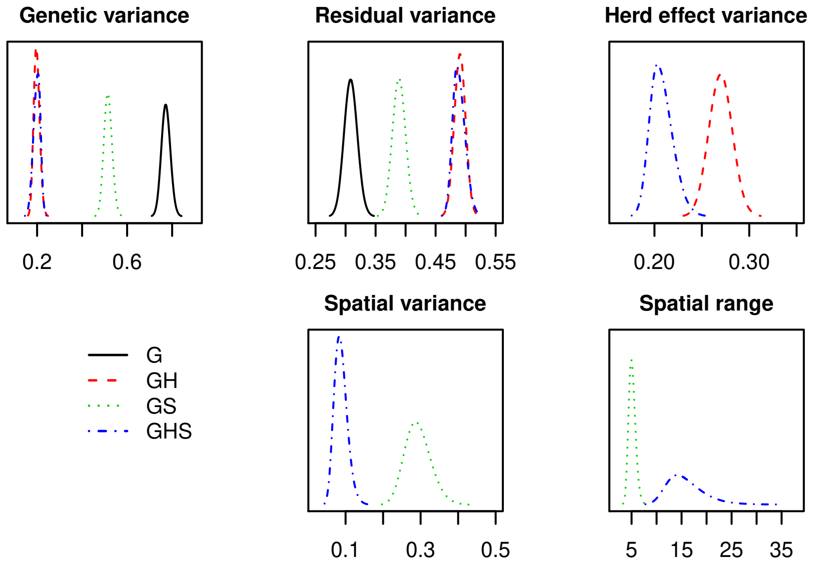


Figure 3: Posterior distributions of hyper-parameters from models G, GH, GS and GHS applied to the full real data

Table 14: DIC for models G, GH, GS and GHS applied to the full real data

| Model | DIC |
|---|---|
| G | 67329 |
| GH | 70964 |
| GS | 70096 |
| GHS | 70929 |

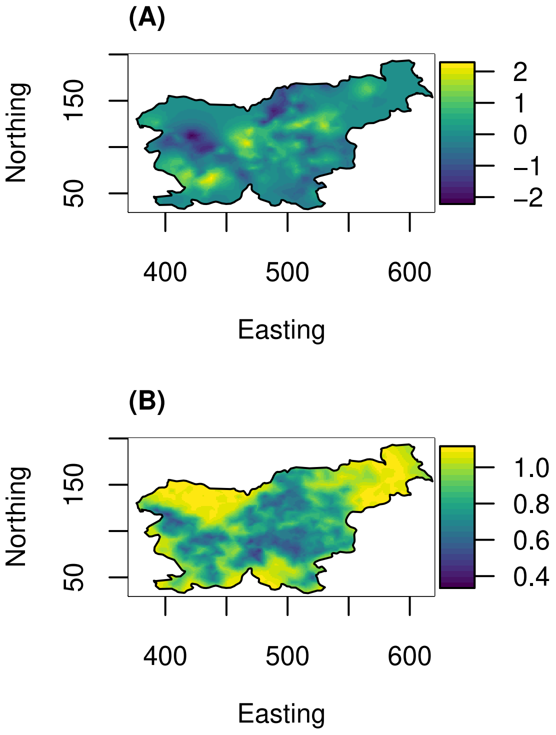


Figure 4: Posterior mean (A) and standard deviation (B) of the estimated spatial effect (in units of spatial standard deviation) from model GHS fitted to the real data - the axis units are in km

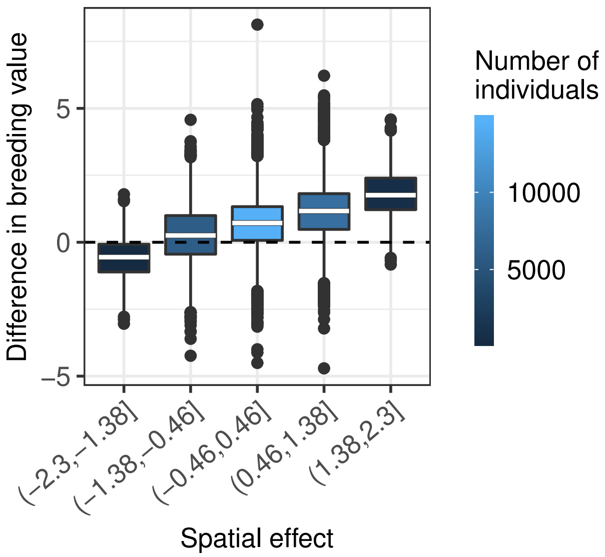


Figure 5: The difference in estimated breeding values (in units of genetic standard deviation) between models GH and GHS by the estimated spatial effect (in units of spatial standard deviation)
